# On the causes, consequences, and avoidance of PCR duplicates: towards a theory of library complexity

**DOI:** 10.1101/2022.10.10.511638

**Authors:** Nicolas C. Rochette, Angel G. Rivera-Colón, Jessica Walsh, Thomas J. Sanger, Shane C. Campbell-Staton, Julian M. Catchen

## Abstract

Library preparation protocols for most sequencing technologies involve PCR amplification of the template DNA, which open the possibility that a given template DNA molecule is sequenced multiple times. Reads arising from this phenomenon, known as PCR duplicates, inflate the cost of sequencing and can jeopardize the reliability of affected experiments. Despite the pervasiveness of this artifact, our understanding of its causes and of its impact on downstream statistical analyses remains essentially empirical. Here, we develop a general quantitative model of amplification distortions in sequencing datasets, which we leverage to investigate the factors controlling the occurrence of PCR duplicates. We show that the PCR duplicate rate is determined by the ratio between library complexity and sequencing depth, and that amplification noise (including in its dependence on the number of PCR cycles) only plays a minor role for this artifact. We confirm our predictions using new and published RAD-seq libraries and provide a method to estimate library complexity and amplification noise in any dataset containing PCR duplicates. We discuss how amplification-related artifacts impact downstream analyses, and in particular genotyping accuracy. The proposed framework unites the numerous observations made on PCR duplicates and will be useful to experimenters of all sequencing technologies where DNA availability is a concern.

## Introduction

The occurrence of polymerase chain reaction (PCR) duplicates is an artifact present in most current sequencing technologies, from whole genome resequencing, to single-cell RNA (Marx, 2017), and to Restriction site-Associated DNA (RAD) sequencing (Table 1). Sequencing library preparation protocols often include a PCR step to improve yield or create molecular species of interest, but this also introduces artifacts (Aird et al., 2011; Kebschull & Zador, 2015). In particular, since these amplified libraries comprise multiple copies of each original template molecule, it becomes possible to independently sequence several reads that correspond to the same template; such reads are known as PCR duplicates (Casbon, Osborne, Brenner, & Lichtenstein, 2011).

**Table 1:** PCR duplicate rates across sequencing technologies.

| Method | PCR Duplicate rate | Reference |
| --- | --- | --- |
| Whole genome sequencing | 1-5% | Ebbert et al. (2016) |
| Exome sequencing | 3-15% | Shigemizu et al. (2015) |
| Exome sequencing | 2-13% | Bonfiglio et al. (2016) |
| Exome sequencing | 15-30% | Borgström et al. (2017) |
| Linked-read DNA sequencing | 11-59% | Meier et al. (2021) |
| Deep targeted DNA sequencing | 3-63% | Smith et al. (2014) |
| Ancient DNA | 2-80% | Gamba et al. (2015) |
| Ancient DNA | 2-42% | Kapp et al. (2021) |
| Single-cell DNA sequencing | 15-25% | Gonzalez-Pena et al. (2021) |
| Single-cell bisulfite sequencing | 2-6% | Farlik et al. (2015) |
| RNA-seq | 1-14% | Fu et al. (2018) |
| RNA-seq | 1-68% | Parekh et al. (2016) |
| Single-cell RNA-seq | 85-95% | Ziegenhain et al. (2017) |
| HiC | 14-64% | Niu et al. (2019) |
| Micro-Capture-C | 80-96% | Hua et al. (2021) |

PCR duplicates create a distorted view of the abundancies of molecules in the original sample, which may bias or degrade downstream statistical analyses (Casbon et al., 2011; DePristo et al., 2011; Fu, Wu, Beane, Zamore, & Weng, 2018; Niu et al., 2019). These concerns have motivated the development of amplification-free methods (Kozarewa et al., 2009; Niu et al., 2019), but such protocols come with substantial technical constraints, particularly with regard to input biological materials. Most methods have instead accepted the artifact as an inherent feature of sequencing data and focus on tracking and removing PCR duplicates bioinformatically (Marx, 2017; Sims, Sudbery, Ilott, Heger, & Ponting, 2014), which can be done either based on read (or read pair) mapping coordinates (DePristo et al., 2011; Li et al., 2009), by tagging template molecules before amplification using unique molecular identifiers (UMIs) (Casbon et al., 2011; Kivioja et al., 2011), or by combining both approaches (Islam et al., 2014; T. Smith, Heger, & Sudbery, 2017).

Although *a posteriori* removal of PCR duplicates has proven effective to mitigate bias, it is not without limitations. This approach can increase the cost of sequencing noticeably—up to severalfold when duplicates represent most of the initial reads (Table 1). In the worst case, after deduplication, coverage may become insufficient for the intended purpose causing experiments to fail. The bioinformatic identification of duplicates is also imperfect. Coordinate-based tracking is unsuitable for experiments in which coverage in some genomic regions is high enough that they become saturated, such as bulk RNA-seq (Fu et al., 2018; Parekh, Ziegenhain, Vieth, Enard, & Hellmann, 2016), ChIP-seq (Tian et al., 2019) or Pool-seq (Kofler, Nolte, & Schlötterer, 2016). UMI-based tracking is confounded by sequencing errors, which must be rigorously accounted for to avoid incomplete deduplication (Islam et al., 2014; Marx, 2017; T. Smith et al., 2017). Lastly, and more fundamentally, removing PCR duplicates is a pragmatic approach that targets the visible consequences of amplification, but distortions can be expected to remain present even after deduplication.

Therefore, developing a better understanding of amplification-related artifacts and gaining control of the rate of PCR duplicates *a priori* by optimizing library preparation procedures remains highly desirable. Despite the prevalence of these artifacts, we still lack a realistic quantitative model for library amplification and the generation of PCR duplicates. As a result, our comprehension of the phenomenon remains highly empirical (Marx, 2017), and there remains some uncertainty and confusion regarding the precise experimental factors that control their occurrence, how these factors interact with one another, and the consequences of PCR duplicates on downstream statistical analyses.

Among the factors believed to have an effect on the PCR duplicate rate, the most important one is library complexity, which has been alternatively defined as the complement of the duplicate rate (Chen et al., 2012), as the information content of a library (Zhang et al., 2015), or as the number of distinct molecular species represented in a sequencing library ((Daley & Smith, 2013); following (Lander & Waterman, 1988)). Insufficient library complexity is frequently given as the probable cause of high duplicate rates (Chen et al., 2012; Marx, 2017; Parekh et al., 2016; E. N. Smith et al., 2014; Tin, Rheindt, Cros, & Mikheyev, 2015), and several studies have demonstrated that the amount of starting biological material used, which presumably correlates with library complexity, had a marked effect on PCR duplicates (Casbon et al., 2011; Fu et al., 2018; Kapp, Green, & Shapiro, 2021; E. N. Smith et al., 2014). A few authors have pointed out that sequencing depth should also be considered (Daley & Smith, 2013; Fu et al., 2018; Marx, 2017; E. N. Smith et al., 2014). Lastly, it is often claimed that PCR duplicate rates depend on the number of PCR amplification cycles that the library was subjected to (K. R. Andrews, Good, Miller, Luikart, & Hohenlohe, 2016; Ebbert et al., 2016; Flanagan & Jones, 2018; Marx, 2017; Orlando et al., 2021; T. Smith et al., 2017; Stuart, Buckberry, & Lister, 2018; Vargas-Landin, Pflüger, & Lister, 2018). Some studies have indeed found this to be the case (Lu, Hofmeister, Vollmers, DuBois, & Schmitz, 2017; Niu et al., 2019; Parekh et al., 2016), but others have concluded that no such relationship existed (Fu et al., 2018; Tin et al., 2015). Thus, several factors relevant to the phenomenon have been identified, but their precise roles remain unclear. A more quantitative understanding of the PCR duplicate artifact would help clarify which methodological alterations are likely to suppress it.

Another question that has been particularly debated in the context of RAD-seq — a reduced-representation sequencing approach that is widely used for population genomics studies of non-model organisms (K. R. Andrews et al., 2016; Catchen et al., 2017; Daley & Smith, 2013) — is the extent to which PCR duplicates affect the reliability of genotyping. While (Flanagan & Jones, 2018; Tin et al., 2015) found it to be an important source of error, (Euclide et al., 2019) did not. In any case, substantial PCR duplicate rates have been reported in some datasets (K. R. Andrews et al., 2016; Davey et al., 2013; Díaz-Arce & Rodríguez-Ezpeleta, 2019; Hoffberg et al., 2016; Schweyen, Rozenberg, & Leese, 2014), which is especially concerning as some popular protocols do not allow the monitoring of this artifact (*e.g.,* double-digest RAD-seq without UMIs). In addition, as sample availability is often a limiting factor in non-model organisms (K. R. Andrews et al., 2016; Peterson, Weber, Kay, Fisher, & Hoekstra, 2012; Tin et al., 2015), better understanding the extent to which observed PCR duplicate patterns result from the use of reduced amounts of DNA would be of great experimental interest. Finally, allelic dropout in RAD-seq has been attributed to restriction site polymorphism (a RAD-seq-specific artifact) (K. R. Andrews et al., 2016), but this may also reflect an incomplete understanding of the consequences of PCR duplicates.

Here, we develop a general quantitative framework able to realistically model amplification-related artifacts in sequencing experiments based on library complexity, sequencing depth, and amplification noise. We provide a method, *Decoratio*, to estimate these factors for any sequencing dataset based on PCR duplicate patterns. We apply this method to new and previously published RAD-seq datasets to demonstrate that amplification artifacts are often an important feature of this sequencing approach and that the model recapitulates the main properties of these experiments. We show that our model reconciles the numerous earlier observations made on PCR duplicate rates, and we discuss how amplification-related artifacts increase variance in downstream analyses, with particular application to genotyping accuracy. Overall, this work furthers our understanding of the properties and consequences of amplification artifacts and will facilitate the optimization and deployment of novel sequencing technologies.

## Methods

### PCR duplicate model

We identify three stages of library preparation and sequencing as being critical to the modeling of PCR duplicates: (1) the pre-amplification pool of template molecules; (2) the pool of amplified molecules; and (3) the pool of molecules that are sampled for sequencing into digital reads (Fig. 1). Accordingly, our model for the occurrence of PCR duplicates comprises two disjoint steps, that respectively connect the first and second, and the second and third of these stages.

**Figure 1:**
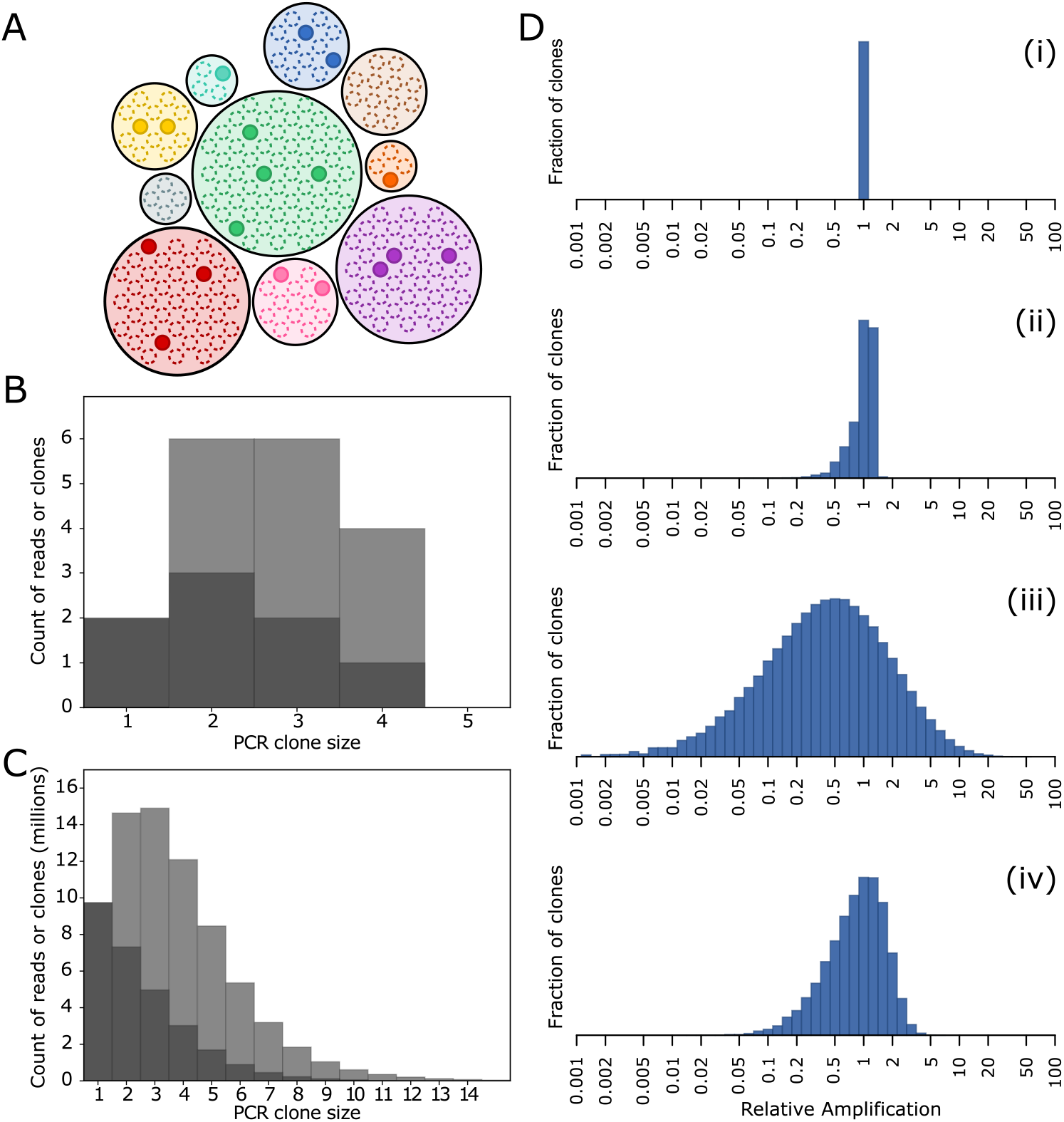
Overview of the PCR duplicate rate model. **(A)** Schematic representation of the processes involved in the occurrence of PCR duplicates. During library preparation, template DNA molecules are amplified, so that for each initial molecule a clone (colored circles) of amplified molecules (dots) is generated. Since PCR is a stochastic and biased process, amplification efficiency is heterogenous across templates and amplification clones vary in size. Subsequently, a small fraction of amplified molecules are randomly sampled for sequencing and become reads (solid dots). Reads that correspond to the same template, *i.e.*, that belong to the same clone, are known as PCR duplicates. Duplicate reads are more likely to be sampled from large clones (*i.e.*, templates whose amplification was most successful) than from small clones. **(B, C)** In practice, what can be observed in sequencing data is the size of clones at the read level (*i.e.*, the number of singletons, of pairs of two duplicate reads, *etc.*). This serves as the main empirical input for the model. The histograms show the distribution of read clone sizes respectively for the schematic in the first panel (B) and for the brown anole dataset (C), either as the number of clones of each size (dark grey) or as the number of reads that belong to clones of a particular size (dark and light grey combined). These views are equivalent by definition, as for instance the number of reads in clones of size two (duplicate pairs) must be twice the number of such clones. Furthermore, as the light grey bars represent redundant reads, the PCR duplicate rate can be calculated from such histograms by taking the ratio of the light grey area over the total area; in this case respectively 56% (18 reads in 8 distinct clones) and 61%. **(D)** Amplification noise is modeled as the distribution of relative amplification factors among templates. Histograms show the amplification factor distributions for (i) a perfect, noiseless amplification; (ii, iii) for the empirically developed amplification model of (Best et al., 2015) with mean amplification efficiency and bias parameters set to “low noise” and “high noise” values, respectively m=0.7, s=0.01, 12 cycles, and m=0.45, s=0.1, 18 cycles; and (iv) for the log-skew normal distribution fitted to the brown anole dataset. These distributions are used to model sequencing data as a mixture of Poisson distributions (see Methods).

In the first step, we model the amplification of template molecules. Namely, given a pool of template molecules, we determine a probability distribution for their individual amplification factors. As the precise method used to determine the distribution does not matter, this approach is highly flexible; we propose two amplification models. First, a distribution of amplification factors can be obtained using forward simulations. For instance, we implemented the PCR model developed empirically by (Best, Oakes, Heather, Shawe-Taylor, & Chain, 2015): starting with one molecule, we apply the amplification model, then record the final number of molecules in the resulting clone (*i.e.*, the amplification factor), and repeat the amplification process independently, one molecule at a time, to obtain an estimate of the distribution of clone sizes. Alternatively, the distribution of amplification factors can be set to some relevant parametric distribution, such as the log-normal distribution.

In the second step, we model the sequencing of reads from the amplified pool of molecules. Crucially, we treat the pool of amplified molecules as infinite, and use the distribution of amplification factors (*i.e.*, of amplification clone sizes) as a statistical description of this pool. This assumption is always reasonable because in practice the amplified pool has to be much larger than the number of reads derived from it—for instance, in the case of Illumina sequencing, only a small fraction of the PCR product is eventually loaded onto a flow cell and bridge-amplified. Under these assumptions, the occurrence of PCR duplicates can be modeled using a Poisson mixture model, as follows.

We refer to the set of duplicate molecules that were amplified from a particular template molecule as an ‘amplification clone’, and to the set of reads that derive from a particular template molecule (*i.e.*, duplicate reads) as a ‘sequencing clone’. Importantly, for a given read dataset, the PCR duplicate rate *r* is a function of the distribution of sequencing clone sizes. For each clone, we count one read as unique and the rest as duplicates, which gives the formula

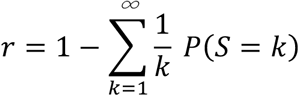

where *P*(*S* = *k*) is the distribution of sequencing clone sizes. The distribution of sequenced clone sizes can be modeled as

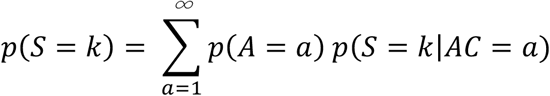

where *P*(*A* = *a*) is the distribution of amplification clone sizes (*i.e.*, the distribution of amplification factors), and

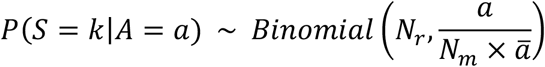

where *N_r_* is the number of read pairs that were sequenced, *N_m_* is the number of template molecules in the pre-amplification pool of DNA, and 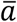 is the mean amplification factor, so that 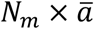 is the number of molecules in the amplified pool of DNA. Assuming *N_r_* is large and no single species dominates in the pool, this can alternatively be written as

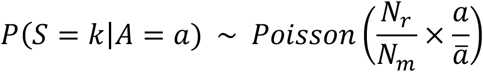

The PCR duplicate rate then depends on two separate parameters: (i) the ratio between the number of sequenced read pairs and the number of pre-amplification template molecules (*i.e.*, the number of unique species in the library, its absolute complexity), which we hereafter refer to as the ‘depth-complexity ratio’, and (ii) the distribution of amplification factors relative to the mean amplification factor, *i.e.*, the noisiness of the amplification (Fig. 1D). It can be noted that the absolute value of the average amplification factor has no effect; this results from our assumption that the pool of amplified molecules is infinite, which, as argued above, is always realistic.

Importantly, it is possible to re-write the depth-complexity ratio in terms of coverage. Specifically, if we consider a particular read (or read pair) length *l_r_* and a set of target genomic regions (*e.g.*, the whole genome) of total length *L_T_* from which the reads derive, then nucleotide-wise coverage can be written as

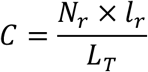

Similarly, we can define molecular coverage—hereafter ‘molecular density’ to avoid confusion around the term coverage—as

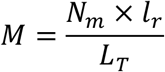

This allows us to re-write the depth-complexity ratio as

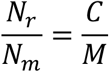

Molecular density defined above is fundamentally a locus-wise measure of library complexity and corresponds to the number of unamplified template molecules that cover an average position of the sequencing target. We note that when coverage is tallied at the nucleotide level, the complexity of a library depends on the read length that is used to sequence it and is maximized when the full length of the molecules is sequenced. This is expected, because the read length influences the number of useful nucleotides within each molecule, and therefore the information content of the library.

### Implementation

In the model described above, the PCR duplicate rate depends on two parameters: the depth-complexity ratio, and the distribution of amplification factors. We provide a method, *Decoratio* (for ‘depth-complexity ratio’), to jointly estimate these two parameters based on the distribution of PCR clone sizes observed in a sequencing dataset. Given that the experimenter already knows the sequencing depth, the depth-complexity ratio also corresponds to an estimate of library complexity.

The program requires two inputs, a distribution of PCR clone sizes and a class of PCR models, and outputs the optimized depth-complexity ratio and PCR model parameters, as well as a plot of the input distribution and fitted model. An example of the expected input, command line call, and outputs of the program is shown in Fig. S1. The distribution of PCR clone sizes should be formatted as a TSV table giving the number of clones of each size, and can be obtained using programs such as *SAMtools-Markdup* (Li et al., 2009), *Picard-MarkDuplicates* (McKenna et al., 2010), *UMItools* (T. Smith et al., 2017), or *Stacks-gstacks* (Rochette, Rivera-Colón, & Catchen, 2019), as described in *Decoratio’s* online manual. If an experiment comprises multiple libraries, we stress that clone size distributions should be derived on a per-library basis, rather than on an aggregated dataset, as the properties of the data are likely to vary across libraries.

For the PCR model, the program currently implements log-normal and log-skew-normal distributions of amplification factors, as well as the empirical “inherited efficiency” model class of (Best et al., 2015) which we described above. The model may be fully specified or only partly so, in which case the program will optimize the model parameters. For most users, using the default log-skew-normal model should work well and the program will fit the standard deviation and skew parameters (Fig. 1). For the (Best et al., 2015) model, the distribution of amplification factors is obtained by forward simulation. Each clone is assigned a duplication probability drawn from a normal distribution, and then amplified by a succession of binomial samplings. In practice, the parameter space is binned to increase computational efficiency, and by default one million simulations are performed. We added a slightly modified variant of this model that uses a Beta distribution instead of the normal distribution (Fig. S2) so as to avoid parametrization issues related to the need to truncate the normal distribution between 0 and 1. The program jointly optimizes the depth-complexity ratio and amplification model by minimizing the sum of squared residuals to the observed distribution of the fraction of reads in each clone size class, using simplicial homology global optimization (SHGO) as implemented in SciPy.

The program is written in Python using NumPy and SciPy and is available from PyPI or from Bitbucket (bitbucket.org/rochette/decoratio) under the GNU GPL v3 license.

### Anolis sagrei RAD libraries

Genomic DNA was extracted from a total of 39 *Anolis sagrei* embryos using the DNeasy® Blood & Tissue Kit (Qiagen) and quantified using a Qubit fluorometer (Life Technologies). Two RAD-seq libraries (Anole-600ng and Anole-30ng) were prepared using a protocol adapted from (Baird et al., 2008; Etter, Preston, Bassham, Cresko, & Johnson, 2011). An average of 280ng of genomic DNA per sample was digested with *SbfI*-HF (NEB) for 60 min at 37°C in a 15μL reaction, followed by heat inactivation for 20 min at 80°C. Custom P1 adapters comprising 10bp barcodes were ligated to each sample using T4 DNA Ligase (Enzymatics, Inc) at room temperature for 20 min, following by heat inactivation for 20 min at 65°C. Samples were then pooled, sheared with a Bioruptor (Diagenode), purified and concentrated with a MinElute Reaction Cleanup Kit (Qiagen). DNA fragments holding inserts in the 150-750bp range were isolated from a 1.5% agarose, 0.5X TBE gel using a MinElute Gel Extraction Kit (Qiagen) and polished with an End Repair/dA-Tailing module (NEB). After subsequent purification, custom P2 Y-adapters were ligated to the DNA fragments at room temperature. The ligated product was purified and quantified using a Qubit Fluorometer with the dsDNA HS assay kit (Invitrogen/Thermo Fisher).

PCR pools comprising 600ng or 30ng of this product, for the Anole-600ng or Anole-30ng libraries respectively, were prepared and divided in 50μL wells in such a way that each well comprised 5ng of DNA, 25μL of 2X Phusion Master Mix (NEB), 2.5μL of 10μM primer mix and up to 22μL H2O. Both libraries were amplified for 18 cycles using the same PCR program. Amplification reactions were pooled, and for each library 300μL was cleaned and concentrated with a PCR Purification Kit (Qiagen), and size-selected as previously described. The final libraries were quantified on an Agilent Bioanalyzer and sequenced 2×100bp on an Illumina HighSeq-4000.

### American Robin RAD libraries

A total of 150 samples of the American robin (*Turdus migratorius)* were collected from central Illinois (Luro et al., *unpublished*). Genomic DNA was extracted from blood samples using the DNeasy® Blood & Tissue Kit (Qiagen) and quantified using a Qubit fluorometer (Life Technologies). A single-digest RAD-seq library was prepared using previously described protocols (Baird et al., 2008; Etter et al., 2011). Genomic DNA was normalized across samples and digested using the *SbfI*-HF enzyme (NEB). Custom P1 adapters containing a 7-bp unique barcodes (Hohenlohe, Bassham, Currey, & Cresko, 2012) were ligated to each individual sample using a T4 ligase (NEB). The 150 robin samples were pooled at equimolar concentrations. Pooled DNA was sheared using a Covaris sonicator and size selected for 300-600bp inserts using Beckman Coulter AMPureXP paramagnetic beads. The sheared DNA was end-repaired and ligated to custom P2 adapters. Before PCR amplification, the pooled DNA of each library was quantified to ensure an exact mass of DNA was added to the reaction. 400ng of template DNA was amplified in a 100μL Phusion HF DNA Polymerase (NEB) reaction (split across four 25μL reactions) for 12 PCR cycles. The final library was sequenced on an Illumina NovaSeq-6000 SP 2×150bp lane.

### Re-analyzed RAD-seq datasets

The stickleback dataset is composed of ten threespine stickleback (*Gasterosteus aculeatus)* samples, sampled from two freshwater and marine populations in Cook Inlet, Alaska (Nelson & Cresko, 2018). The samples were prepared using a single-digest RAD-seq library (Baird et al., 2008; Etter et al., 2011) using the *PstI* restriction enzyme and sequenced using 2×250bp paired-end reads (NCBI-SRA project PRJNA429207).

The warbler dataset consists of 241 yellow warbler (*Setophaga petechia*) individuals sampled from 18 populations across North America (Bay et al., 2018). Samples were processed in three separate libraries: 95 samples from 8 populations in *Warbler-1*, 89 samples from 9 populations in *Warbler-2*, and 68 samples from 14 populations in *Warbler-3.* All three libraries were prepared using the SbfI-based BestRAD protocol (Ali et al., 2016) and sequenced with 2×100bp paired-end reads (NCBI-SRA project PRJNA421926).

The penguin dataset contains sequencing of 66 of Emperor penguin (*Aptenodytes forsteri*) samples collected across four populations in Antarctica (Cristofari et al., 2016). The samples were prepared across three separate libraries: *Penguin-3* with 30 samples across 2 populations, *Penguin-4* with 24 samples from 1 population, and *Penguin-81* with 12 samples from 1 population (R. Cristofari, personal communication). All libraries were processed into single-digest RAD-seq libraries (Baird et al., 2008; Etter et al., 2011), digested with *SbfI*, and sequenced with 2×100bp reads (NCBI-SRA project PRJNA308448).

### Bioinformatic analyses of RAD datasets

All RAD libraries were analyzed separately using the *Stacks* v.2.5 pipeline (Rochette et al., 2019). For newly sequenced libraries (Anole and Robin), the raw sequence data was first demultiplexed based on samples’ RAD barcodes using the `process_radtags` program with flags `--paired, --rescue` and `--clean-quality`. For all libraries, RAD loci were assembled *de novo* using the `denovo_map` script in paired-end mode with mismatch parameters `M` and `n` kept equal and set to respectively 5, 7, 9, 5, and 5 for the Anole, Robin, Stickleback, Warbler, and Penguin datasets; this is primarily to account for differences in read length. We identified and removed PCR duplicates (`--rm-pcr-duplicates`) and selected only loci present in a majority of samples (flag `-X “gstacks: --dbg-min-loc-spls”` was set to half the number of samples in each library). Finally, the populations program was used to export variants genotyped in at least 50% of samples (`-r 0.5`) in VCF format.

### qPCR quantification of library complexity

To obtain an empirical measure of the complexity of the robin library, the amount of template DNA was quantified using qPCR. This template represents DNA from the RAD library after P2 ligation, prior to amplification by PCR. The template DNA of the library was amplified alongside the final RAD library, which is cleaned and quantified prior to sequencing. The known concentration of the final library was used to then calculate an absolute concentration of template in the pre-amplified library. To create a standard curve, five 1:10 sequential dilutions of the 0.1nM control library were prepared and amplified in triplicate, alongside a negative control. Similarly, the library template was diluted in two sequential 1:10 dilutions and amplified in triplicate. qPCR reactions were prepared using the KAPA Library Quantification Kit (KAPA Biosystems). While the KAPA kit by default uses primers compatible with Illumina’s P5 and P7 oligo sequences, the complimentary sequence for these primers is not present in single-digest RAD-seq libraries until it is reconstructed during PCR amplification (Baird et al., 2008; Etter et al., 2011). Instead, we performed qPCR amplification using the primers designed for sdRAD library enrichment PCR (forward primer: 5’- AATGATACGGCGACCACCGAGATCTACACTCTTTCCCTACACGACGCTCTTCCGATC*T-3’, reverse primer: 5’- CAAGCAGAAGACGGCATACG*A -3’) (Baird et al., 2008; Etter et al., 2011; Hohenlohe et al., 2012). For the reaction, the forward and reverse primers were combined into a single 10μM primer mix. Each 20 μL reaction consisted of 10.4μL of KAPA SYBR FAST mix with ROX, 2μL of standard Primer Mix, 3.6μL of PCR-grade water, and 4 μL of DNA. The qPCR reactions were run on a QuantStudioTM 3 Real-Time PCR System (Applied Biosystems) using the default ΔΔCT protocol. This protocol consists of an initial denaturation step of 1 minute at 95°C, 35 cycles of a 95°C 30 seconds denaturation followed by a 60°C annealing/extension/data acquisition for 45 seconds, and a 65-95°C melting curve analysis.

The CT of a given sample was obtained by calculating the average across its replicates. The concentration of the library template in nM was obtained by regressing its average CT value against the standard curve obtained for all the known control.

## Results

### PCR duplicate rates in RAD-seq datasets

To assess the general occurrence of PCR duplicates in RAD-seq studies, we reanalyzed a series of new and published datasets. Specifically, we generated two single-digest RAD (sdRAD) (Baird et al., 2008) paired-end datasets, respectively comprising 39 brown anole (*A. sagrei*) individuals and 150 American robin (*T. migratorius*) individuals. We also considered the paired-end datasets from (Cristofari et al., 2016), (Nelson & Cresko, 2018), and (Bay et al., 2018), which respectively use sdRAD in the Emperor Penguin (3 libraries; Penguin-3, Penguin-4 and Penguin-81), sdRAD in the Threespine Stickleback (1 library), and bestRAD (Ali et al., 2016) in the Yellow Warbler (3 libraries; Warbler-1 to Warbler-3). We did not include any libraries based on ddRAD (Peterson et al., 2012) as this protocol does not allow the tracking of PCR duplicates (but see Discussion).

Overall, we found significant PCR duplicate levels in all analyzed datasets. Mean per-library duplicate rates ranged from 21% for the Robin library to 95% for Penguin-3 (Fig. 2; Table 2). Unsurprisingly, because a 95% duplicate rate corresponds to a sequencing efficiency of just 5%, non-redundant coverage (*i.e.*, coverage after removing PCR duplicates) was very low for the samples of the Penguin-3 library, ranging from 1.6 to 4.3x. However, even the better libraries were subject to an appreciable sequencing efficiency loss. The bestRAD protocol, yielding three libraries with low duplicate rates, tended to perform better than the sdRAD protocol, which yielded libraries with both high and low duplicate rates, although our sample of datasets may be too small for generalization.

**Figure 2:**
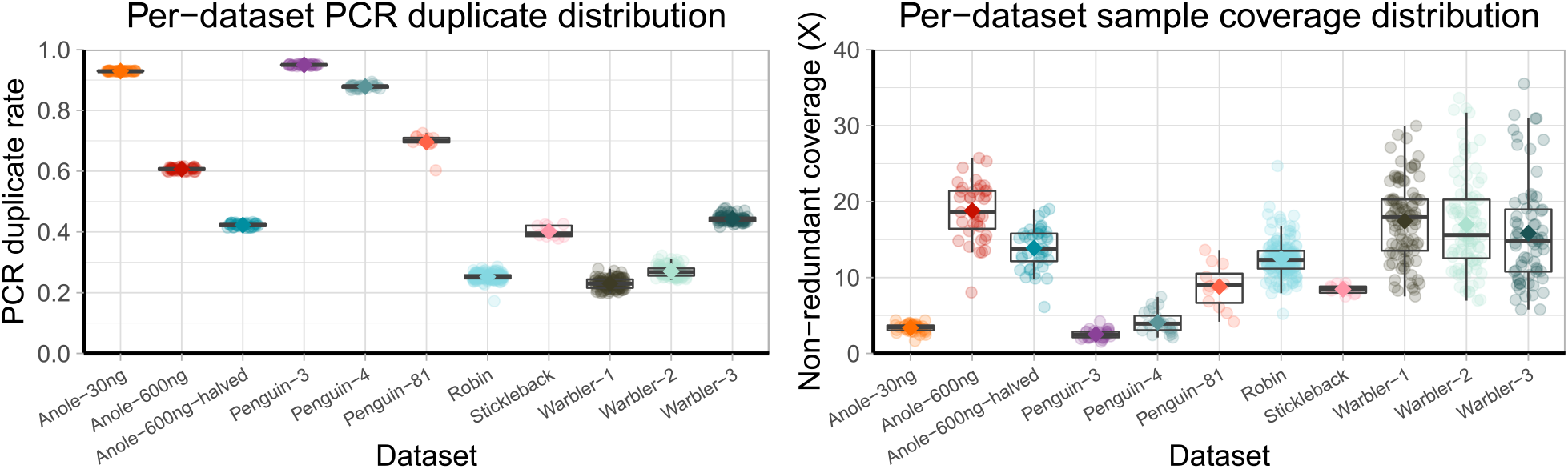
PCR duplicate rates in reanalyzed sdRAD datasets. **(A)** Distribution of PCR duplicates for all samples in each dataset. Colored diamonds show the mean per-library duplicate rate. The PCR duplicate rate is consistent across all samples within a molecular library. For datasets comprised of multiple libraries, the duplicate rate can vary among the different libraries. **(B)** Non-redundant coverage distribution for all samples in each dataset. Colored diamonds show the mean non-redundant coverage. In contrast to PCR duplicate rate, coverage is highly variable within these libraries.

**Table 2:** PCR duplicate rates and library complexities in new and reanalyzed RAD-seq datasets.

| Library | Anole 600 | Anole 30 | Robin | Stickle-back | Penguin 3 | Penguin 4 | Penguin 81 | Warbler 1 | Warbler 2 | Warbler 3 |
| --- | --- | --- | --- | --- | --- | --- | --- | --- | --- | --- |
| Raw library coverage (total) | 1899x | 1895x | 2452x | 144x | 1532x | 820x | 351x | 2218x | 2133x | 2006x |
| Samples in library | 39 | 39 | 150 | 10 | 30 | 24 | 12 | 95 | 89 | 68 |
| PCR duplicate rate | 61 ± 1% | 93 ± 1% | 21 ± 1% | 40 ± 2% | 95 ± 0.2% | 88 ± 1% | 70 ± 3% | 23 ± 2% | 27 ± 2% | 44 ± 1% |
| Depth-complexity ratio | 1.93 | 14.2 | 0.29 | 0.87 | 19.9 | 8.16 | 1.85 | 0.38 | 0.46 | 0.85 |
| Library density (total) | 984x | 133 | 8455x | 166x | 77x | 101x | 190x | 5837x | 4637x | 2360x |
| Library density (per-sample mean) | 25x | 3.4x | 56x | 17x | 2.6x | 4.2x | 16x | 61x | 52x | 35x |
| Mean non-redundant coverage | 19.1 ± 3.8x | 3.4 ± 0.6x | 12.8 ± 2.4x | 8.6 ± 0.6x | 2.6 ± 0.6x | 4.2 ± 1.5x | 8.9 ± 2.8x | 17.9 ± 5.3x | 17.3 ± 6.4x | 16.3 ± 7.2x |

Perhaps most importantly for the purposes of this work, it appeared very clearly that the relevant level at which to look at PCR duplicates was the library, rather than the individual sample or the study. Indeed, PCR duplicate rates varied greatly across libraries within each dataset (for datasets comprising several libraries) but were highly consistent across samples within each molecular library (Fig. 2; Table 2). Remarkably, quality differences between individual DNA samples are likely present in at least some of the tested libraries but did not appear to impact PCR duplicate rates.

These observations prompted us to investigate the causes underlying the occurrence of PCR duplicates, the potential detrimental effects of their presence, and the steps that should be taken to reduce these rates.

### A realistic model for PCR duplicates

We present a quantitative model that captures the steps of library preparation and sequencing that are critical with regard to PCR duplicates. Briefly, this model comprises two steps. First, we model how the relative molecule abundancies within a DNA library are distorted by PCR amplification. Specifically, for a chosen PCR model, we derive a distribution of the relative amplification factors across molecules within a library (Fig. 1C). Second, we derive expected patterns of PCR duplicates (*i.e.*, the distribution of PCR clone sizes in the sequencing data; Fig. 1B) by modeling the stochastic sequencing process - accounting for sequencing depth, the complexity of the library, and amplification distortions – using a compound Poisson model. In practice, the model uses two parameters: a PCR model, and the ratio between the number of sequenced reads and the number of unique molecules in the library, which we refer to as the depth-complexity ratio. These parameters are fitted to individual datasets by matching the predicted distribution of PCR clone sizes to the observed one.

Using this method, we were able to reproduce the clone size distributions that were observed experimentally in the datasets introduced above (Fig. 3A-B, S3). Accounting for amplification led to considerably better fits than using a noiseless, simple Poisson model, especially in datasets that have more pronounced amounts of PCR duplicates (*e.g.*, Fig. 3A). Such datasets provide more information on the underlying true distribution of amplification factors, whereas in datasets with low duplicate rates, most reads are singletons and are uninformative in this regard. In addition, even if the noiseless model may appear to be a reasonable fit for datasets with fewer duplicates (*e.g.*, Fig. 3B), ignoring amplification noise still led to skewed library complexity estimates Table S1. For instance, for the 22%-duplicate Robin dataset, the true library complexity was likely more than 60% larger than the estimate based on a noiseless model.

**Figure 3:**
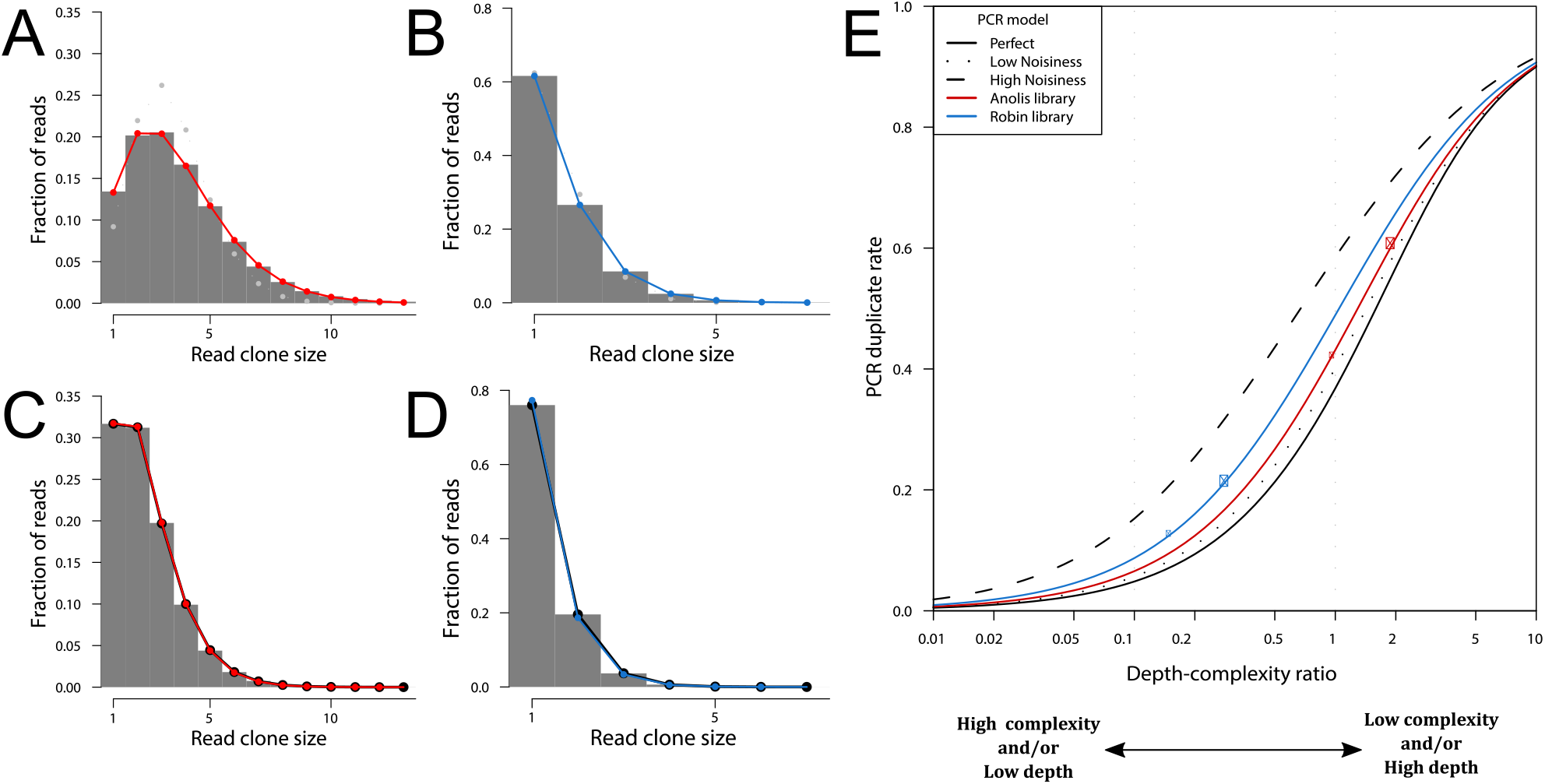
Expected relationship between the depth-complexity ratio and the PCR duplicate rate. **(A, B)** Histogram of the read clone sizes observed in the Anole-600ng and Robin datasets, respectively, overlaid with the fitted depth-complexity ratio and amplification noise model (solid red and blue lines respectively) or a noiseless amplification model (Poisson distribution; dotted gray lines). **(C, D)** Same as (A) and (B) respectively, but for reanalyses of each dataset after randomly discarding half the reads (twofold downsampling). The red (respectively blue) line shows the distribution predicted when halving the depth-complexity ratio in the model fitted on the full dataset (*i.e.*, the colored lines in panels A and C respectively). In both panels, the thick black line shows the model fitted to the downsampled dataset. **(E)** Expected relationship between the PCR duplicate rate, the sequencing depth, the library complexity, and the PCR itself. The solid, dotted, dashed and red amplification noise models are the same as in (Fig. 1D), and the blue model is the best fit to the American robin dataset. Large colored dots mark the observed PCR duplicate rates and estimated depth-complexity ratios for the brown anole (red) and American robin (red) RAD-seq datasets, respectively, and small dots the values observed when considering only half the reads in those datasets, as shown in panels (A-D). PCR duplicate rates are determined primarily by the ratio between the sequencing depth and the complexity of the library, and modulated by the noisiness of the model used for the amplification. The curves shown in this panel may also be considered from the perspective of sequencing saturation (Fig. S6).

Remarkably, our model also captures one striking property of empirical RAD-seq libraries: that all samples within a library had almost identical PCR duplicate rates, regardless of any differences in coverage or DNA quality that may exist among them (Fig. 2). The model suggests this happens because the primary determinant of the rate of PCR duplicates is the depth-complexity ratio. This ratio is expected to be identical for all samples within a library, because for each sample both the number of reads and the number of (active) template molecules are measures of the relative abundancy of that sample in the library. For instance, a sample representing one percent of a library’s template molecules can be expected to later receive one percent of the total sequencing coverage for this library, so that the depth-complexity ratio of that sample will be equal to the global depth-complexity ratio of the library.

Nevertheless, we note that there remained a little within-library variation of the PCR duplicate rate (Fig. 2), which attests to the presence of sample-specific effects. Some, but not all of this residual variation could be explained by differences in library representation (41% and 71% of the variance for the Anolis and Robin libraries respectively; Fig. S4). Conceivably, differences in DNA quality among samples may lead to different responses during PCR amplification or sequencing. For instance, the fragment size distribution may vary across samples, and this could result in differences in mean amplification factor and/or sequencing efficiency.

### PCR duplicate rates vary predictably with coverage

While differences in coverage among the samples in a library due to unequal representation do not prevent them from exhibiting the same PCR duplicate rate, the dependence of PCR duplicate rates on the depth-complexity ratio nevertheless implies a more general dependence between coverage and PCR duplicates. Specifically, for a given library, the ratio’s denominator (*i.e.*, complexity) is fixed once the library has been prepared, whereas the numerator depends on the sequencing effort that is subsequently applied. Consequently, for a given library, increasing the sequencing effort should increase the PCR duplicate rate by a predictable amount.

We tested this prediction using the *A. sagrei* dataset by down-sampling the original reads to reduce the sequencing coverage by half, with the expectation that the PCR duplicate rate would decrease accordingly. The original dataset had a raw coverage of 49x and an observed PCR duplicate rate of 61% (Table 2). The fitted model predicted that the depth-complexity ratio in this experiment was 1.93, and that halving the coverage should decrease the PCR duplicate rate to 42%. After processing the down-sampled dataset, a PCR duplicate rate of 42% was observed, exactly matching the prediction. At a finer level, the change in the shape of the distribution of PCR clone sizes was also predicted precisely (Fig. 3A-D). We conclude that our model captures essential properties of the PCR duplicate phenomenon and has predictive value.

### PCR duplicate rates depend primarily on the complexity of a library

Next, we summarize the predictions of the model regarding the behavior of PCR duplicate rates under a range of scenarios. As previously described, the model relies on two parameters: the depth-complexity ratio and a PCR model. The results obtained by varying these parameters are presented in (Fig. 3E).

Importantly, while the PCR duplicate rate depends on both depth and complexity, it is only sensitive to the value of the ratio between the two, regardless of their respective absolute values. In the model, this property derives from assumptions on the sequencing sampling process, but it also holds for real data (see above results regarding the uniformity of duplicate rates within libraries). We thus only investigate variations of the depth-complexity ratio and did not assess the effects of coverage and complexity individually. Similarly, with regard to the PCR model parameter, the duplicate rate only depends on the distribution of relative amplification factors (Fig. 1D), regardless of the mean absolute amplification factor or of the precise mechanism generating the spread of amplification factors. What matters especially is the overall ‘noisiness’ of the PCR—whether all molecules are amplified equally or, on the contrary, whether some molecular species become much more abundant than others. For this reason, the results presented here, derived using the class of PCR models developed empirically by (Best et al., 2015), should be robust to PCR model choice, and specifically should generally hold for any PCR model producing an approximately lognormal distribution of amplification factors. We note that the amplification noisiness range assessed here is relevant for typical library amplification reactions; some specialized amplification techniques, such as multiple displacement amplification, are much noisier (Gawad, Koh, & Quake, 2016) and may thus fall outside of the considered range.

Our central observation is that depth-complexity ratios much less than one always yield low duplicate rates, and that depth-complexity ratios greater than one always yield high duplicate rates. The duplicate rate is significantly influenced by the PCR model only when the depth-complexity ratio is between one tenth and one (Fig. 3E). Nevertheless, the duplicate rate is always in the 5-15% range if the depth-complexity ratio is 0.1, or in the 35-60% range if that ratio is 1, regardless of the noisiness of the PCR.

Thus, we find that the PCR duplicate rate depends primarily on the depth-complexity ratio, and only marginally on the PCR model. From an experimental perspective, it should be noted that the sequencing coverage is set according to experimental needs (*e.g.*, to 30x) whereas complexity is an intrinsic and difficult to control property of the prepared library. Consequently, a high depth-complexity ratio typically occurs because the library has a molecular density that is too low with regards to experimental needs. Thus, the above result can most simply be understood as: high PCR duplicate rates occur when pre-amplification library complexity is insufficient, largely independently of the PCR protocol being used.

### qPCR measurement of the molecular density of RAD-seq libraries suggests in silico estimates are realistic

It is notable that the library complexities measured above (Table 2) do not match the values expected from the physical mass of DNA used to prepare the libraries. For instance, the PCR reaction of the 150-sample Robin library used a total of 400 ng of DNA. Given that the weight of one haploid American robin genome (*i.e.*, its C-value) is 1.39×10^-3^ ng (C. B. Andrews, Mackenzie, & Gregory, 2009), the DNA mass used is equivalent to 288,000 genome copies, so that the library could be expected to have a density of 288,000x (or 1,918x per sample, on average). In contrast, our modeling of PCR duplicate patterns in the resulting sequencing data suggests a total density of 8,500x (57x per sample). Thus, the diversity of molecules represented in sequence reads was 34 times less than the mass-based expectation.

To explain this discrepancy, we hypothesized that only a small fraction of the molecules present could actually be amplified and subsequently sequenced, while most molecules would be degraded or otherwise inert for the purpose of amplification and sequencing (Meyer et al., 2008). To test this experimentally, we used qPCR to quantify amplifiable DNA in the pre-PCR molecular library. We measured a template single-stranded DNA concentration of 0.192 nmol/L (Fig. S5), which implies that the 40 μL used during library preparation corresponded to 7.68×10^-6^ nmol, *i.e.*, 4.62 billion, single-stranded template molecules. Owing to the design of adapters in the RAD-seq protocol used here, these molecules each corresponded to independent double stranded molecules, *i.e.*, contributed fully to library complexity. Additionally, the density of the library was bioinformatically measured over the 89,162 loci found in more than half the samples, and these loci collectively represented 57.5% of the reads in the library (with the rest of read pairs corresponding to repetitive RAD loci, to RAD loci that are only found in one or a few individuals, or to genomic DNA not flanking with a restriction site). As this proportion must also hold in the library at the molecule level, we distributed 57.5% of 4.62×10^9^ molecules (*i.e.*, 2.66×10^9^ molecules) over 89,162 loci, so that we obtained a qPCR-based library density estimate of 29,800x.

Thus, the experimental, qPCR-based library density estimate was much lower (9.7 times less) than the estimate based on the C-value (288,000x) (Fig. 4; Table 3). This confirmed that most of the DNA used for this amplification reaction was indeed inert, and that estimates of library complexity based on DNA mass alone may be inflated by more than an order of magnitude. Nevertheless, we note that the qPCR-based estimate remained 3.5 times higher than the *in silico* one; we cannot presently resolve this residual discrepancy.

**Figure 4:**
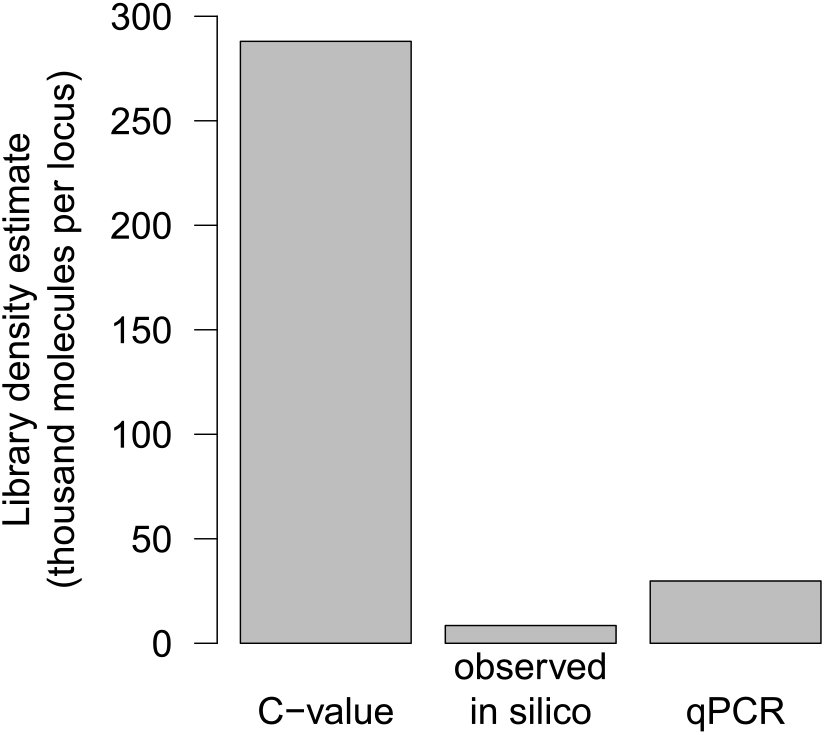
Library complexity is much smaller than suggested by DNA mass. Library complexity for the American robin RAD-seq library, measured as the total molecular density for 150 pooled samples, obtained using proxies available at different stages of the experiment. Calculating library complexity by dividing the mass of DNA used for amplification by the C-value of the organism vastly over-estimates the value actually observed in downstream in silico analyses. Quantifying this same DNA using qPCR highlights that most molecules are indeed not amplifiable and therefore do not contribute to library complexity, owing to sample preservation and partial yield at earlier experimental steps.

**Table 3:** Library and bioinformatic statistics needed to contextualize an experiment’s PCR duplicate rate. For sequencing approaches other than RAD-seq, the number of RAD loci should be substituted with a relevant measure of the size of the genomic target, and most libraries will only include one sample. The American Robin C-value was sourced from (C. B. Andrews et al., 2009) and the Anole C-value (1.97) was calculated assuming a genome size of 1.93 Gbp (Geneva et al., 2021) and a DNA weight of 1.023 pg/Gbp (Doležel, Bartoš, Voglmayr, & Greilhuber, 2003).

| Library | Anole-600 | Robin |
| --- | --- | --- |
| DNA amplified (ng) | 600 | 400 |
| DNA amplified (C-value equivalents) | 305,000 | 288,000 |
| Total aligned read pairs | 72,942,759 | 189,422,046 |
| PCR duplicate rate | 61% | 21% |
| Number of filtered RAD loci | 43,906 | 89,161 |
| Samples in library | 39 | 150 |
| Mean non-redundant coverage | 19.1x | 12.8x |

### Increasing library complexity reduces PCR duplicate rates experimentally

Finally, the comparison between our two *Anolis sagrei* RAD-seq libraries, Anole-600ng and Anole-30ng, illustrates the experimental importance of library complexity. These two libraries were prepared from the same 39 DNA samples using protocols that were exactly identical except for the volume used at the amplification step, respectively 6,000 and 300μL, which corresponded to 600 and 30ng of template DNA. The yield of the smaller reaction was not technically limiting so that subsequent steps could be performed identically, and both libraries were then sequenced at the same depth of 49x.

As predicted, the observed library density was much higher for the Anole-600ng library than for the Anole-30ng one, respectively at 984x and 133x. Accordingly, the PCR duplicate rates were respectively 61% and 93% (for a sequencing depth of 49x), resulting non-redundant coverages of 19.1x and 3.4x per sample, on average (Table 2). This adds to previously published evidence (see Discussion) to demonstrate the critical role of the total amount of DNA used for PCR amplification, independently of all other experimental factors, in determining library complexity and PCR duplicate rates.

In addition, further inspection of these two datasets highlighted that library density was an intrinsic and separate property of these datasets, rather than merely a parameter fitted so that the model replicates PCR duplicate patterns. Since a low per-sample library density indicates that only a few molecules are stochastically sampled and amplified at each locus, we can expect that the makeup of the library itself will have a pronounced stochastic component, so that the resulting read data will be more variable than expected based on the randomness of the sequencing process alone (Fig. 5A; and see Discussion).

**Figure 5:**
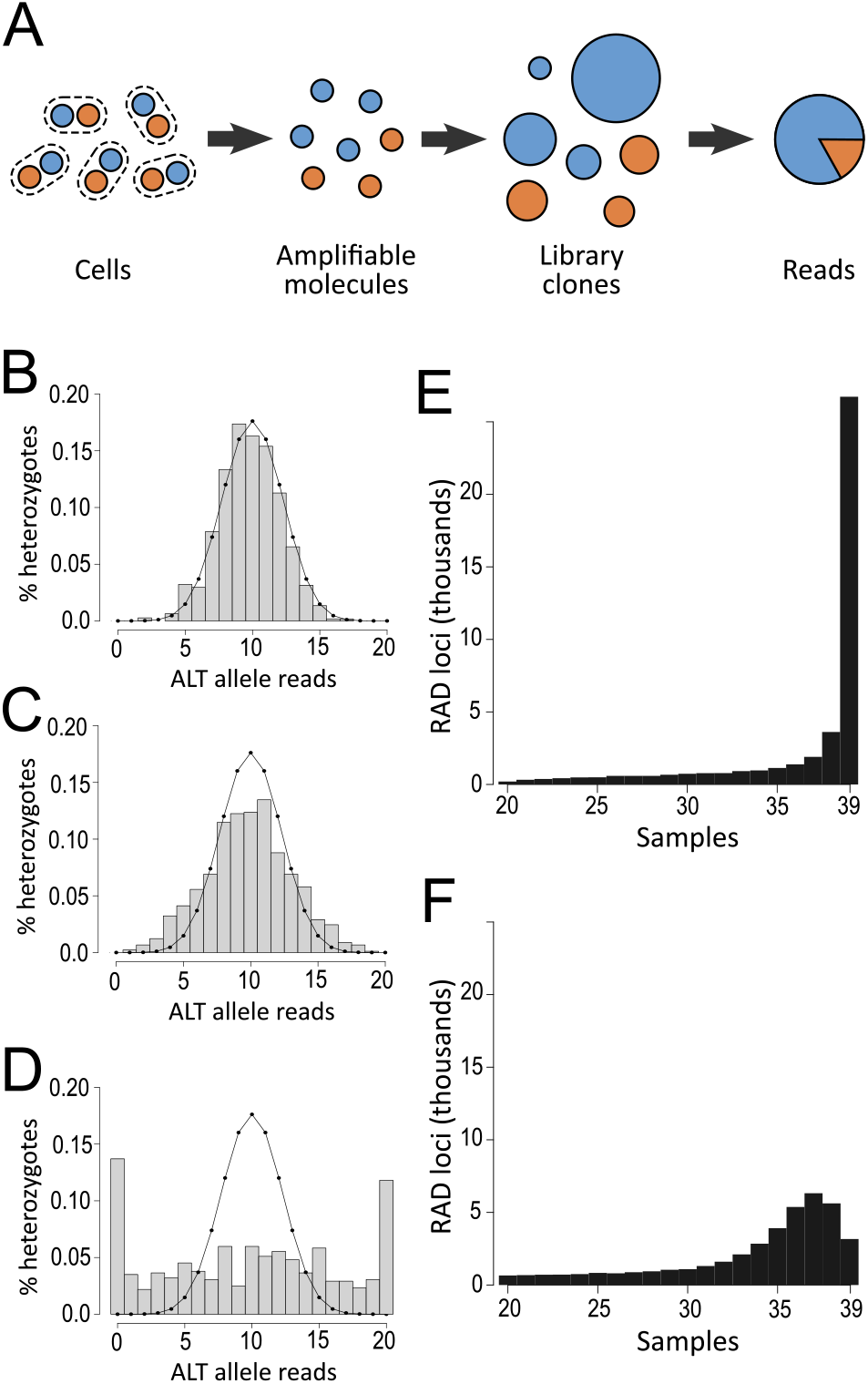
Consequences of low library complexity on genotyping. **(A)** Due to molecule sampling and noisy amplification, the alleles at a heterozygote locus may already be unequally represented in the library by the time it is sequenced, causing sequence data to be overdispersed compared to the binomial distribution assumed by genotyping models. This artifact is expected to be particularly serious at low library complexities and can be mitigated by removing PCR duplicates (see Discussion). **(B-D)** Observed allelic counts at heterozygous sites of anole individual E35 that have a total depth of 20 reads, respectively in the Anole-600ng dataset with (B) or without (C) PCR duplicate removal, and in the low-complexity Anole-30ng dataset without PCR duplicate removal (D). The solid line shows the binomial distribution. Peaks at x=0 and x=20 in (D) are due to allelic dropout, where one allele is entirely absent from the library. Heterozygous sites were annotated based on the Anole-600ng dataset with PCR duplicate removal. **(E, F)** Histograms showing apparent locus sharing among the 39 individuals of the Anole dataset, respectively for the Anole-600ng and Anole-30ng libraries. The latter, Anole-30ng library, exhibits “locus dropout” because most individuals have such a low molecular density that at any locus several individuals typically fail to sample even a single amplifiable molecule.

This over-dispersion is first apparent for coverage patterns at heterozygous sites. Considering the sequencing process alone, the number of reads observed for either allele should follow a binomial distribution. The Anole-600ng data approximately conforms to this expectation, especially when the effects of amplification stochasticity are mitigated by removing PCR duplicates Fig. 5B-C. In the Anole-30ng dataset, however, the number of observations for the two alleles deviate much more from equal representation, and in many cases one allele simply isn’t observed Fig. 5D. The latter is explained by allelic dropout: given that the per-sample density of the library is very low, at only 3.4 molecules per locus or 1.7 molecules per allele, on average, frequently no molecules will be sampled for a particular allele.

Second, at such low densities, there is also a chance that neither of the two alleles of a locus will be sampled, so that an entire locus may be stochastically dropped. While in RAD-seq analyses it is normal that loci aren’t perfectly shared across individuals due to polymorphisms in restriction sites, comparing the extent of locus sharing in the Anole-600ng and Anole-30ng datasets (Fig. 5E-F) makes it clear that most of the variation in locus composition across individuals in the Anole-30ng data is caused by stochastic locus dropout due to low library density rather than genetic polymorphism.

## Discussion

### Determinants of PCR duplicate occurrence

PCR duplicates are a pervasive sequencing artifact and a wealth of empirical observations on their occurrence have been reported. However, this knowledge accumulation has often happened as a by-product of the development and validation of new molecular protocols, and the lack of a theoretical framework within which to connect individual results has not allowed a complete understanding of the artifact’s causes and of its seriousness in specific applications (Marx, 2017). We fill this gap by introducing a mechanistic and quantitative model for the occurrence of PCR duplicates. This allows us to unify earlier results, to further clarify the effects of various experimental factors, and to draw expectations about the statistical properties of duplicate-containing datasets.

Our first observation is that the PCR duplicate rate fundamentally depends on the ratio between sequencing depth and library complexity (Fig. 3). In particular, we stress the symmetry between depth and complexity, and that it is imperative to consider both when comparing PCR duplicate rates across experiments. The idea that PCR duplicates occur when a library is sequenced in excess relative to its complexity has been discussed in earlier works (Daley & Smith, 2013; Fu et al., 2018; Rao et al., 2014; T. Smith et al., 2017), and in principle could be extrapolated following (Lander & Waterman, 1988). However, these studies did not consider amplification artifacts or focused on a specific problem or application. For instance, the method of (Daley & Smith, 2013) proposed to estimate library complexity based on the species-saturation approach of (Efron & Thisted, 1976), which ultimately amounts to modeling amplification noise in the same way as presented here. However, as the authors’ focus is solely on library complexity, this factor is then eliminated through non-parametric approximations and its effects are not discussed further.

The central role of library complexity and sequencing depth is experimentally supported by the RAD-seq-based results presented here, as well as by observations from earlier studies. The amount of material used as input for library preparation has been shown to strongly impact duplicate rates in RNA-seq (Fu et al., 2018) and ancient DNA (Kapp et al., 2021). Very high duplicate rates are also apparent in whole genome resequencing libraries prepared from minute amounts of DNA (*e.g.*, Bruinsma et al., 2018), and there is evidence that this parameter is also important in ATAC-seq and Hi-C (see below). As for the effect of sequencing depth, it is implicit in experiments where sequencing is performed to saturation (Daley & Smith, 2013; Farlik et al., 2015; Niu et al., 2019; Ziegenhain et al., 2017) and should generally be immediately apparent in any dataset containing significant levels of duplicates if the read data is down-sampled (Fig. 3).

Second, since our model explicitly accounts for amplification, we were able to explore its behavior across a range of amplification parameters. We find that PCR itself only plays a minor role in the rate of occurrence of PCR duplicates. Although a greater variance in relative amplification factors across templates (*i.e.*, a “noisier” amplification) does elevate PCR duplicate rates, it only does so within a narrow range determined primarily by the depth-complexity ratio (Fig. 3C).

This conclusion is in contrast with the frequent assertion that PCR duplicates are due to an excessive number of PCR cycles (Ebbert et al., 2016; Marx, 2017; Orlando et al., 2021; T. Smith et al., 2017; Stuart et al., 2018; Tin et al., 2015; Vargas-Landin et al., 2018). The absence of a major effect of PCR is nevertheless in agreement with published experimental results. In particular, (Fu et al., 2018) specifically tested this interaction in RNA-seq and found none. Similarly, (Tin et al., 2015) observed no change in RAD-seq duplicate rates when increasing the number of cycles. And while several studies have reported a strong effect of the number of cycles, in RNA-seq (Parekh et al., 2016), ATAC-seq (Lu et al., 2017), RAD-seq (Díaz-Arce & Rodríguez-Ezpeleta, 2019), and Hi-C (Niu et al., 2019), in each case the amount of starting material for the protocol was made to vary concurrently, so that either factor could be responsible for the observed differences. In fact, (Fu et al., 2018) also made this observation, but subsequently found that amplification was not the causal factor. Thus, given theoretical expectations and other empirical results in this direction, these studies can more parsimoniously be interpreted as further evidence that library complexity plays a pivotal role in a wide range of sequencing approaches.

Although from an experimental perspective it makes sense to think jointly of the amount of material available and the number of amplification cycles to perform, confounding their effects hinders the formulation of clear and effective recommendations to improve experiments compromised by PCR duplicates. In particular, numerous authors have suggested that limiting the number of cycles was critical to minimize duplicates (Marx, 2017; Orlando et al., 2021; Stuart et al., 2018). Using fewer cycles should indeed lead to fewer PCR duplicates because it forces experimenters to use more starting material to achieve necessary yields. However, the reverse is not true; if enough starting material is used, excess amplification should not appreciably impact the resulting duplicate rate. Thus, we find that with regard to PCR duplicates, experimenters should not be excessively concerned with the amplification protocol and should instead focus primarily on increasing the amount of (active) starting material for the amplification, even if this is the hardest thing to do as it may call for substantial alterations to the protocol (*e.g.*, Kapp et al., 2021). Nevertheless, general optimization of the amplification is still advisable because a higher efficiency is directly associated to a lower amplification noise (see below).

### Measuring library complexity

Although library complexity and sequencing depth have exactly opposite roles in determining the PCR duplicate rate, this symmetry is only mathematical as in practice these two factors come with very different constraints. Sequencing depth is an experimental choice and is set by the experimenter to a level called for by the intended application (*e.g.*, upwards of 30x for single-individual germline genotyping (Sims et al., 2014), 1x for low-coverage population genotyping (Lou, Jacobs, Wilder, & Therkildsen, 2021), or 10-30M reads for RNA-seq in mammals (Stark, Grzelak, & Hadfield, 2019)) and can be adjusted *a posteriori* by sequencing again if required. Library complexity, in contrast, is established permanently during library preparation so that the only way around insufficient library complexity is to prepare new libraries (Rao et al., 2014). In this context, it is critical for experimenters to achieve an appropriate complexity at the time of library preparation. A complexity at least one order of magnitude higher than the required sequencing depth is ideal for most approaches, although an interesting exception is linked-read approaches (Meier et al., 2021; Zheng et al., 2016) where a high sequencing saturation is necessary to sample multiple fragments associated with each tag, so that library complexity should instead be matched to the intended sequencing depth (Weisenfeld, Kumar, Shah, Church, & Jaffe, 2017).

The experimental number which is most relevant to library complexity is the number of initial molecules that effectively undergo amplification. Once a library has been amplified, each unique molecular species is present in many copies and is therefore unlikely to be lost, so that the complexity then remains essentially constant at later steps of the library preparation protocol. In other words, library complexity is fundamentally determined by the amount and quality of DNA that is input into the PCR. Consequently, reducing PCR duplicate levels primarily amounts to amplifying more template DNA. However, it is important to stress that only a fraction of a pool of DNA is usually amplifiable, so that mass is a poor proxy for the amount of useful DNA (Gansauge & Meyer, 2013; Kapp et al., 2021; Meyer et al., 2008) and the complexity of libraries prepared using the same input mass but from samples differing in quality or using distinct protocols may greatly differ. A more direct estimate of library complexity can be obtained by qPCR quantification of the library before it is amplified, as this directly measures the concentration of active molecules. This assay also informs on the percentage yield of the protocol that was used (Gansauge & Meyer, 2013; Kapp et al., 2021), which ultimately is what determines how complex a library derived from a given finite sample can be.

### Measuring library complexity as a molecular density

As noted above, the minimum acceptable library complexity—measured as an absolute number of distinct molecules—may vary by several orders of magnitude depending on the intended application. This definition of library complexity is thus only meaningful to compare closely related experiments, rather than in a general sense, making it difficult to establish guidelines. Here, we argue that measuring library complexity in terms of the number of molecules per locus will often be more informative than the above “absolute” library complexity. In the context of sequencing approaches aiming for a homogenous coverage (*e.g.*, whole-genome, exome, bisulfide, or RAD-seq), scaling absolute library complexity by the size of the genomic regions considered leads to a measure that can be directly compared to coverage, and which we refer to as the molecular density of a library. As this measure considers the sequencing target, it becomes possible to provide quantitative recommendations. For instance, for single-individual genotyping at 30x coverage, library molecular density should ideally be greater than 300x, assuming a one-order-of-magnitude margin. This target value applies regardless of whether the sequencing target is a whole genome or an exome, or if the organism considered has a genome size substantially different from that of mammals, as is the case for many species of medical, agricultural, or ecological interest (*e.g., Drosophila melanogaster, Danio rerio, Arabidopsis thaliana*).

In addition, the molecular density of a library is more relevant than absolute complexity both when considering amplification-related statistical error patterns in downstream computational analyses (see below) and when evaluating experimental yield. The latter is because it is directly related to units that are used for experimental inputs, such as number of cells (*e.g.*, (Brind’Amour et al., 2015; Butler, 2015; Lu et al., 2017)), genome equivalents (*e.g.*, (Lander & Waterman, 1988; Zheng et al., 2016); mass expressed in units of the C-value), or template DNA units ((Taberlet et al., 1996); equal to two genome equivalents). The relationship with yield is most obvious when the input is measured in number of cells: if one starts an experiment with 1,000 diploid cells, comprising 2,000 genomes copies, the maximum number of unique molecules that may be sequenced for any genomic region is also 2,000. Molecular density measures how many molecules per locus (on average across the genome) are present in the final library, and therefore represents the overall yield. Taking this logic to the limit highlights that molecular density is also closely related to the ‘genome coverage’ yield statistic used in single-cell whole-genome sequencing experiments (Daley & Smith, 2014; Zhang et al., 2015): especially, for haploid cells sequenced to saturation, the two become identical.

For the same reason, comparable library densities may be achieved from a given amount of input sample for whole-genome, exome, or RAD-seq, even though the absolute library complexities will vary by orders of magnitude (with no effect on the success of the experiment, as proportionately less molecules are needed to describe an exome than a whole genome). And remarkably, this rationale also applies to heterogenous-coverage approaches. For instance, when performing ChIP-seq on a histone mark for which few peaks exist, for a given number of input cells the resulting yield and absolute library complexity can be expected to be lower than for a more frequent mark, but this does not necessarily imply that the local molecular density (which determines the statistical properties of the data, see below) will be lower at the rare-mark peaks, because the molecules in the library are spread out over fewer loci.

Thus, when examining the yield of a library preparation protocol, we recommend expressing input sample amounts in number of cells or in genome equivalents (*i.e.*, multiples of the C-value mass), and to measure library complexity as a molecular density rather than as an absolute number of unique molecular species.

### Modeling amplification-related overdispersion

Amplification-related artifacts cause increased technical variance (Casbon et al., 2011), which has been demonstrated for instance in RNA-seq (Castel, Levy-Moonshine, Mohammadi, Banks, & Lappalainen, 2015; Fu et al., 2018; Parekh et al., 2016), single-cell RNA-seq (Grün, Kester, & van Oudenaarden, 2014; Islam et al., 2014; Ziegenhain et al., 2017), iCLIP (T. Smith et al., 2017), or Hi-C (Niu et al., 2019). These artifacts have also been reported to cause genotyping errors (K. R. Andrews & Luikart, 2014; Bresadola, Link, Buerkle, Lexer, & Wegmann, 2020; Díaz-Arce & Rodríguez-Ezpeleta, 2019; Taberlet et al., 1996; Tin et al., 2015). The conceptual framework presented here provides insights on the mechanisms that give rise to these error patterns.

Essentially, underlying our model is the view that technical variance in sequencing data is the result of three successive sampling steps: (i) the acquisition of actually amplifiable molecules from the biological sample, (ii) the noisy amplification of these molecules, and (iii) the selection of amplified molecules for sequencing to generate reads (Fig. 1A, 5A). In many methods of sequence data analysis, variance is modelled primarily after read depth, thus only the third step above is accounted for. This may lead to inaccurate variance estimations, particularly if the magnitude of amplification-related artifacts varies among libraries in the considered dataset, or worse, this may cause methods that don’t estimate residual variance (*e.g.*, genotyping, see below) to produce unreliable results by failing to acknowledge the overdispersion of the data. How mis-specified a model is for a particular dataset depends on the relative scales of the variances introduced at each of the three steps, and especially whether the third step–sequencing–is actually the predominant source of variance in the dataset.

Both the first and the third steps are Poisson processes, with means respectively equal to the molecular density of the library and to the sequencing coverage. The second step, amplification, is a complex process, but generally its variance (*i.e.*, the unequal amplification of templates) can be partitioned in two components: systematic bias and stochasticity (Best et al., 2015; Kebschull & Zador, 2015). Systematic bias is caused by differences in amplification efficiency among templates (*e.g.*, due to differences in molecule length) and simply increases geometrically with the number of cycles. In contrast, stochasticity, which can represent most of the variance (Kebschull & Zador, 2015), is due to the partial efficiency of amplification at each cycle (some molecules are amplified and some are not) which is magnified by the exponential nature of PCR. This stochasticity is strongest during the first few cycles when clone sizes are still small, then subsides as clones grow and clone-wise efficiency becomes predictable (Kebschull & Zador, 2015). The mechanisms underlying both variance components impact amplification efficiency, so that amplification variance is minimal when efficiency is high.

The PCR duplicate phenomenon intersects this three-step variance model in several ways. First, as discussed above, an absence of PCR duplicates is an indication that the molecular density of the library is much greater than the sequencing coverage. This directly implies that the variance of the first step’s Poisson sampling (of amplifiable DNA molecules) is negligible in comparison with that of the third step’s Poisson sampling (of reads). In addition, the presence of a large number of molecules for each locus causes stochastic amplification effects to average out, so that the relative variance of the second step is also reduced. Thus, if the PCR duplicate rate is negligible, the first and second steps can be neglected. On the contrary, a high PCR duplicate rate implies that sequencing coverage is comparable to or greater than the molecular density of the library. In this case, the variances associated with molecule sampling and noisy amplification can be comparable to or exceed the variance associated with sequencing itself, leading to increased technical variance and model mis-specification.

In addition, while PCR duplicates can indicate serious underlying amplification artifacts, at the same time they offer the opportunity to monitor and reduce these artifacts. Tracking PCR duplicates allows one to unwind the three-step process *a posteriori.* In the extreme case, sequencing a library to saturation while removing duplicates should lead to an apparent sequencing coverage equal to the library’s molecular density and a complete reduction of amplification noise, essentially simplifying the entire sampling process to only its first step—the sampling of amplifiable molecules. This saturation effect has been observed in single-cell RNAseq (Islam et al., 2014).

In datasets with intermediate duplication levels, the effects of removing PCR duplicates is less straightforward. However, if we assume no systematic differences in amplification efficiency between templates, and considering that removing PCR duplicates is equivalent to keeping track of which amplification clones have received at least one read, then it becomes possible to merge all three steps into a single Poisson process with mean equal to the non-redundant coverage. This re-emergence of the Poisson distribution after PCR duplicate removal suggests that the common practice of using bioinformatic methods designed around the sequencing process alone to process de-duplicated datasets is reasonable.

### Effects of PCR duplicates on genotyping

We will now focus on genotyping, but we note that the mechanistic framework described here should also be useful to better understand technical noise in other sequencing technologies. For instance, in RNA-seq, it represents a possible explanation for the nonlinear relationships between technical noise, input starting material, and gene abundancy (Brennecke et al., 2013) — technical variance at a particular gene should be determined by the local molecular density, which is proportional to both the absolute complexity of the library and to the gene’s expression level. In single-cell RNA-seq, it could inform how the occurrence of a major technical noise feature, gene dropout (Kiselev, Andrews, & Hemberg, 2019), is influenced by the complexities of the sub-libraries for each cell, the noisiness of the amplification, and the relative sequencing effort.

Amplification artifacts have long been known to be a source of error for genotyping (Pompanon, Bonin, Bellemain, & Taberlet, 2005; Taberlet et al., 1996), and it is standard practice to remove PCR duplicates before genotyping. As discussed above, this approach should suppress model mis-specification and yield reliable genotypes, even in cases where initial PCR duplicates rates are high—provided that duplication clones can be identified accurately and that the resulting coverage is sufficient.

However, several recent studies have interrogated the degree to which retaining PCR duplicates impacted the accuracy of genotype calls (K. R. Andrews & Luikart, 2014; Bresadola et al., 2020; Díaz-Arce & Rodríguez-Ezpeleta, 2019; Euclide et al., 2019; Flanagan & Jones, 2018; Tin et al., 2015) as certain RAD-seq approaches do not permit the identification and removal of duplicates (reviewed in (K. R. Andrews et al., 2016)). Surprisingly, while some studies have confirmed the intuitive expectation that PCR duplicates negatively impact genotyping (Bresadola et al., 2020; Díaz-Arce & Rodríguez-Ezpeleta, 2019; Euclide et al., 2019; Tin et al., 2015) have recently reported convincing evidence that genotyping could be reliable even in datasets containing substantial duplicate rates.

Our three-step model allows us to make specific predictions on the effects of amplification-related artifacts on genotyping and to reconcile these apparently contradictory results. The model suggests that amplification-related artifacts impact genotype calls by causing overdispersion of the allelic ratios observed at heterozygous sites. Genotyping models are based on the binomial distribution, with modifications to account for sequencing errors (DePristo et al., 2011; Li, 2011; Maruki & Lynch, 2017; Rochette et al., 2019). This, however, assumes that the allelic ratio is balanced in the final library being sequenced, which can be incorrect in amplified libraries (Fig. 5A). Genotyping models thus underestimate the likelihood of imbalanced allelic ratios when the underlying genotype is a heterozygote. This leads to bias against heterozygotes, which manifests itself either as a loss of power to call heterozygotes, or in the worst case as calling a heterozygote site as homozygote.

Remarkably, how incorrect genotyping becomes if PCR duplicates are not removed doesn’t just depend on the duplicate rate (itself determined by the depth-complexity ratio), but also on the absolute values of sequencing coverage and library density. This is because calling discrete genotypes involves a strong threshold effect. Indeed, a heterozygote site will be called correctly as long as the heterozygote likelihood is significantly larger than the homozygote one. This of course depends on the model, but also on the data itself: if coverage and density are large enough that several reads are consistently observed for both alleles, homozygote likelihoods will be consistently small, so that correct calls can be made regardless of model mis-specification.

In practice, the key statistic is the non-redundant coverage—the apparent coverage after duplicates have been removed, which depends on both sequencing coverage and library density. If the non-redundant coverage is high enough for reliable genotype calls to be made, for instance greater than 20x, it can be expected that calls made without removing duplicates should also be reasonable. This seemingly was the case in the data of (Euclide et al., 2019), which may explain why they found, in apparent contradiction with other studies, that removing PCR duplicates had little influence on genotyping. The argument also applies to the results of (Ebbert et al., 2016), whose data featured a PCR duplicate rate of just 2%.

In contrast, if the non-redundant coverage is too low for reliable genotype calling, attempting to genotype samples without removing PCR duplicates will lead to important biases. The strong overdispersion of the allelic ratios observed at heterozygous sites implies that genotyping models underestimate the chance that all the sampled reads come from the same allele and may thus confidently (but wrongly) call homozygous genotypes at sites that are actually heterozygous. This is essentially a theory of the occurrence of allelic dropout (Broquet & Petit, 2004; Taberlet et al., 1996) in high-throughput sequence data and explains the main error patterns reported in the literature, namely miscalled heterozygotes and deflated heterozygosity (Bresadola et al., 2020; Díaz-Arce & Rodríguez-Ezpeleta, 2019; Flanagan & Jones, 2018; Tin et al., 2015).

This allelic dropout phenomenon is most obvious and most pronounced in low complexity libraries, that have a density smaller than 10x. For instance, if a heterozygote locus is represented by just three molecules, chances are high that all three will represent the same allele. If twenty reads are then sequenced and presented to a genotyping model, this model will confidently infer a homozygote, because it sees a single allele in a relatively large sample size. Assuming the three molecules have each been sequenced at least once, removing PCR duplicates would leave three reads, and the model would correctly conclude that such a small sample size comprises too little information to make a reliable genotype call.

We conclude that genotypes reported without information about the PCR duplicate rate may be unreliable, and therefore that it is crucial to monitor PCR duplicate rates in genotyping experiments. We confirm that removing PCR duplicates is a simple and efficient strategy to mitigate the detrimental effects of amplification-related artifacts on genotyping, and that it comes at a very small cost in power since the information that is discarded is mostly redundant. Although datasets with relatively high PCR duplicate rates can sometimes produce acceptable genotypes even without removing duplicates (Euclide et al., 2019), this happens, perhaps frustratingly, precisely when the non-redundant coverage is high and avoiding this filter presents little appeal. Nevertheless, removing PCR duplicates is preferable even in these cases, as full-coverage genotypes should still be expected to suffer from lesser artifacts such as biased genotyping confidence scores.

### Implications for RAD-seq studies

Our results and those of others (Schweyen et al., 2014; Tin et al., 2015) demonstrate that PCR duplicates are present at substantial levels in both single-digest and double-digest RAD-seq experiments (Table 1). Given that genotyping quality is primarily determined by non-redundant coverage (see above), and consequently that genotype calling is unreliable in experiments where PCR duplicates are not monitored, controlling for PCR duplicates in RAD-seq experiments is critical. We thus advocate the systematic use of protocols that allow their identification, *i.e.*, either single-digest protocols (Ali et al., 2016; Baird et al., 2008) combined with paired-end sequencing, or double-digest protocols combined with UMIs (Hoffberg et al., 2016; Schweyen et al., 2014; Tin et al., 2015). This is essential for demographic analyses, as they tend to be more sensitive to genotyping errors than population structure analyses, and are sensitive in particular to biased estimates of heterozygosity and/or of the number of singleton alleles.

In addition, these results highlight that PCR duplicate rates vary extensively across experiments. This variation must be partly due to differences in library complexity, but we stress that sequencing depth also plays an important role (Fig. 3). In comparing RAD-seq libraries, the measure of choice for library complexity should be library density, which determines both the maximum achievable non-redundant coverage and the cost-efficiency of sequencing (*i.e.*, the number of reads required to achieve a given non-redundant coverage). Importantly, most studies do not report enough information to estimate the complexities of their libraries. We therefore encourage researchers to report all the technical values relevant to understanding the occurrence of PCR duplicates in RAD-seq, namely: the number of samples that were pooled in each library, the mass of DNA that was amplified, the number of RAD loci kept in the analysis, the per-library number of (possibly paired) reads aligning to these loci, the per-library average PCR duplicate rate, and the per-library average number of non-redundant reads per locus.

Nevertheless, the widespread occurrence of PCR duplicates suggests that RAD-seq experiments would typically benefit from libraries with higher densities. As discussed above, this involves amplifying greater quantities of template DNA whenever possible. How much DNA should ideally be amplified depends on the mass of the haploid genome of the study organism (*i.e.*, its C-value), on the quality of the input DNA and percentage yield of the protocol, as well as—crucially for pooled libraries such as those typically seen in RAD-seq— on the number of individuals in the library. Indeed, pooling does not change the per-sample library complexity requirements for genotyping. Pooled libraries therefore require handling considerable amounts of DNA (at least before amplification) to achieve similar per-sample complexities. For instance, amplifying 1 microgram of genomic DNA should not be out of the ordinary for a 100-sample library, as this represents 10 nanograms per sample. This logic is not directly applicable to bestRAD (Ali et al., 2016) because amplification is performed on purified (restriction-site digested, ligated and bead-captured) DNA rather than on genomic DNA; however, all numbers above can instead be applied to the genomic DNA input to the bead-capture step.

Importantly, insufficient library densities can explain allelic dropout, which is another error pattern that has been extensively discussed in the context of RAD-seq, often in combination with concern about restriction site polymorphism (K. R. Andrews et al., 2016; Davey et al., 2011; Gautier et al., 2013). However, as underlined by the heterozygosity deficit caused by amplification-related artifacts, allelic dropout can also result from the purely stochastic non-sampling of one of the two alleles of a heterozygote, which is particularly likely to happen in libraries that have a low molecular density. This is in fact the mechanism that the original definition of allelic dropout refers to (Broquet & Petit, 2004; Taberlet et al., 1996).

Moreover, population genetics simulations suggest that restriction site polymorphism should have little consequences unless genetic diversity is very high (Gautier et al., 2013; Rivera-Colón, Rochette, & Catchen, 2021). In contrast, insufficient library density can create dramatic allelic dropout, and can even result in locus dropout for densities so low that it becomes likely that neither allele is sampled (Fig. 5). Thus, we suggest that in the absence of PCR duplicate tracking, evidence for allelic dropout in a dataset should be considered indicative of a low library density (and of a high duplicate rate) rather than of restriction site polymorphism. Finally, we note that stochastic allelic dropout is a non-issue in datasets where duplicates have been removed, as using non-redundant reads prompts the genotyping models to account for the phenomenon.

## Conclusions

We propose a realistic and predictive model for the distortions that amplification introduces in sequencing libraries. This establishes a conceptual framework within which the numerous observations made on PCR duplicates can be united, the causes of the artifact can be quantified, and their consequences on downstream analyses understood. In particular, we find that:

- PCR duplicate rates are determined mainly by the ratio between sequencing depth and library complexity. Nevertheless, as most experiments target a particular sequencing depth, amplifying a large enough DNA pool is critical to maintain a low depth-complexity ratio and limit PCR duplicates; thus, the amount of active DNA input into the amplification reaction during library preparation is key.
- In most cases, it is advantageous to measure library complexity not as an absolute number of molecules, but as a molecular coverage—molecular density—as this statistic is more transferable between experiments, more relatable to experimental yield, and more directly interpretable into expected error patterns in statistical analyses.
- Because PCR duplicate rates depend on both library complexity and sequencing depth, the reporting and interpretation of PCR duplicate rates should always be accompanied by depth considerations.
- PCR amplification is stochastically uneven, and this has an effect on the statistical properties of libraries, including a minor role in determining duplicate rates and a lowering of the apparent library complexity. However, the number of cycles can be expected to only have a marginal effect on duplicate rates, which is in contrast with a widespread conception, but in agreement with the experimental results that exist in the literature. It is important to maximize per-cycle amplification efficiency, as this reduces stochasticity and bias.
- Our framework can be leveraged to better understand the error patterns that amplification noisiness creates in downstream statistical analyses. We demonstrate this for the case of genotyping, which makes us able to propose recommendations to improve the effectiveness of RAD-seq experiments.

## Author contributions

NR, ARC, SC and JC designed experiments. NR and ARC performed experiments. NR drafted the manuscript. ARC and JC edited the manuscript. JW prepared the *Anolis* sequencing libraries. TS provided and prepared the *Anolis* samples. SC and JC provided funding. All authors approved the final manuscript.

## Acknowledgements

The authors would like to thank Mark E. Hauber and Alec B. Luro for access to the robin RAD-seq dataset and Sarai H. Stuart for assistance with the qPCR.

## Glossary

*Absolute library complexity*: The total number of distinct molecular species in a library; *i.e.*, the number of PCR clones in the library; *i.e.*, *the* number of template molecules that were effectively amplified.
*Genomic target*: The set of genomic regions targeted during library preparation, such as the whole genome, exome, or the regions flanking a particular restriction site or associated with a particular histone mark.
*Molecular density*: A locus-wise measure of library complexity, equal to the average number of distinct template molecules covering loci within the genomic target of a library, or a specific subset thereof (*e.g.*, a particular basepair, or a particular exon).
*Non-redundant coverage*: Coverage after PCR duplicate removal.
*PCR clone*: The set of all the molecules and reads that descend, through PCR amplification, from a particular template molecule.
*PCR duplicates*: A set of reads that belong to the same PCR clone are said to be PCR duplicates. This is a symmetrical relationship: all the reads of a clone are duplicates of one another.
*PCR duplicate rate*: The fraction of reads removed during PCR duplicate removal. This is equal to one minus the ratio of the number of non-redundant reads over the total number of reads. Also known as the clonality rate.
*PCR duplicate removal*: This consists in discarding all but one read for each PCR clone. This removal is typically performed at random to avoid introducing systematic bias. Also known as deduplication, decloning, or clonal reduction.
*Template molecule*: A DNA molecule directly extracted from the biological sample and originally present as a single identifiable molecule (though possibly double-stranded) before the library is amplified.

## Supplementary Tables & Figures

**Supplementary Table S1:**
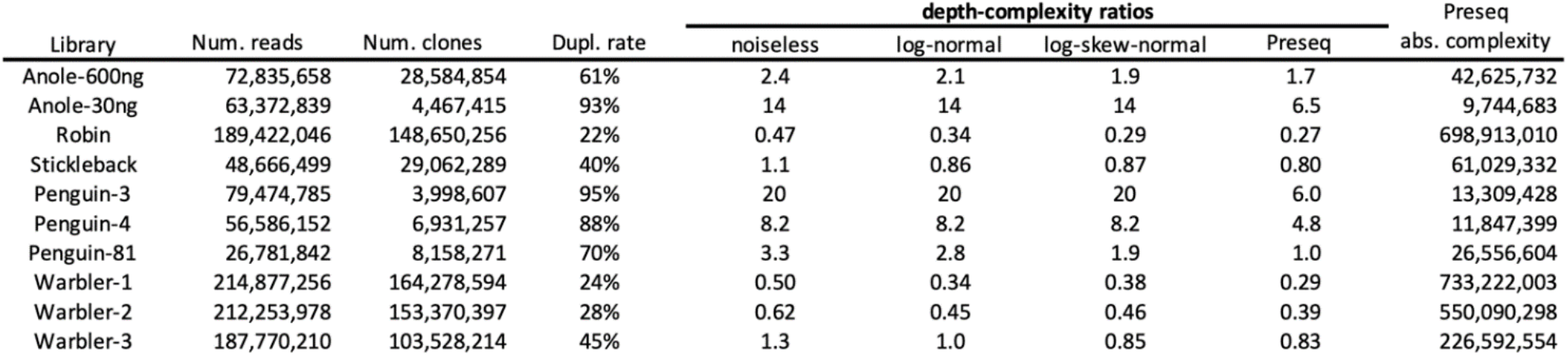
Variability of complexity estimates across methods.

Depth-complexity ratios for all analyzed datasets were estimated with our method using three increasingly complex amplification model families: noiseless, log-normal, or log-skew-normal. The fits for these models are shown in [Supplementary Figure S3]. Additionally, library complexity estimates derived using the Preseq method of (Daley & Smith, 2013) are reported. Preseq absolute library complexity estimates were obtained by taking the number of clones (“distinct reads”) predicted for a depth tending towards infinity (in practice 1,000 billion reads). The corresponding depth-complexity ratio is then calculated by dividing the number of (actually sequenced) reads by the absolute complexity.

Estimates are overall consistent across models, with a tendency that the noiseless model estimates the lowest library complexities (*i.e.*, highest depth-complexity ratios), followed by the log-normal model then the log-skew-normal model. This is expected as for a given data set and duplicate rate, higher complexity estimates must be associated with a higher predicted abundancy of rarely detected, poorly amplified clones. For the same reason, the complexity estimate derived using a log-skew-normal model may be lower than that derived using a log-normal one (for Stickleback, Warbler-2, and especially for Warbler-1) when the optimized amplification factor distribution is positively skewed (*e.g.*, [Supplementary Figure S3], panel H).

Preseq estimates are highly discordant for datasets that have a high duplicate rate. This likely results from an artifact affecting Preseq when analyzing datasets that have already been sequenced at a high saturation—which it wasn’t designed to do. Indeed, it can be seen in [Supplementary Figure S3] that even in duplicate-rich datasets, for which the modal clone size is high, the size-1-clone class tends to be more populated than the size-2-clone class. This does not make physical sense unless the true distribution of amplification factors was extremely bimodal, which would be very unexpected for such genotyping libraries. Instead, this pattern is likely to result from the occasional incomplete clustering of large clones—the accurate and exhaustive annotation of large duplicate clones being delicate (T. Smith et al., 2017). The methodological design of Preseq emphasizes the singleton class, also it is likely to be particularly affected by such signals. Consequently, it may assume the presence in the library of numerous low-abundancy clones and report an inflated absolute library complexity estimate.

In less saturated datasets, however, Preseq estimates are concordant with those of our method, with a tendency to estimate slightly higher complexities. The largest difference is seen for Warbler-1. Again, this discrepancy may be due to an artifact. As mentioned above, for this dataset our method estimates a positively skewed distribution of amplification factors. This reflects the presence, in the input, of a noticeable amount of very large clones: for this dataset 1% of reads belong to clones of size 30 or more (this is in sharp contrast with *e.g.*, the Robin dataset, where such clones do not exist at all). The existence of such large clones suggests differences in amplification efficiency that are simply unrealistic, also the presence of an experimental and/or bioinformatic artifact is likely.

**Supplementary Figure S1:**
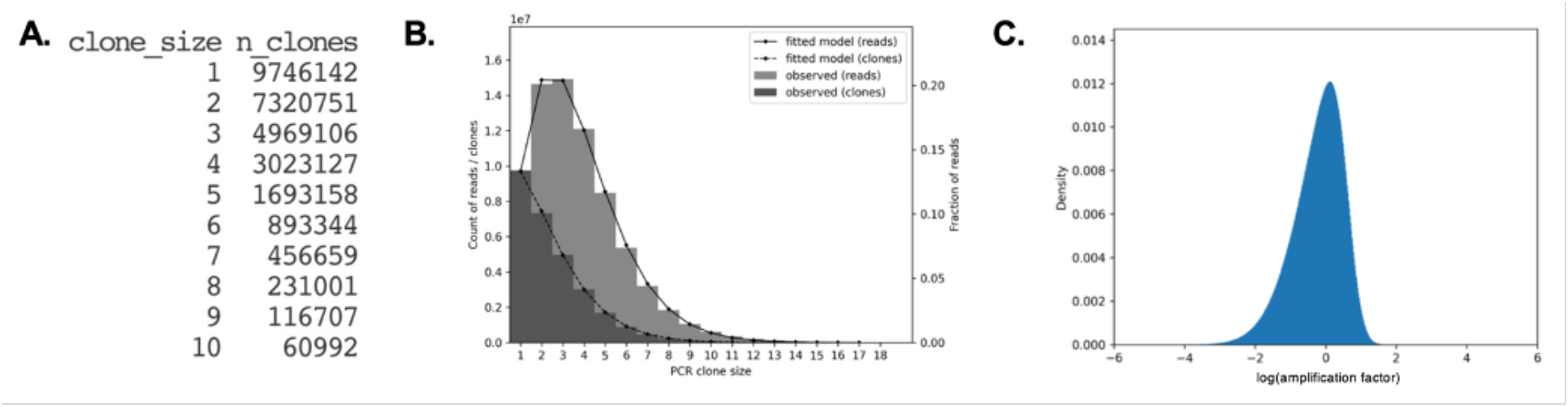
Program inputs & outputs. (A) The input for the method takes the form of a TSV table giving the frequency of each PCR clone size (such that the total number of reads for each clone size is equal to the number of clones of that size multiplied by the clone size, *i.e.*, the product of the two columns), as shown here (truncated at 10 for display). The program may then be called as simply “`python3 -m decoratio clone_size_frequencies.tsv`” if the user wants to use the default (and recommended) log-skew-normal amplification model. (B) The output consists in the optimized numerical values for the depth-complexity ratio and amplification model, plus a PNG plot of the input histogram overlaid with the fitted model. This plot allows for a quick visual check of the coherence of the input and fit of the model. As in main text Figure 1B, dark grey represents the number of clones of each size and combining the dark and light grey gives the number of reads that belong to clones of a particular size. (C) A plot of the distribution of amplification factors corresponding to the optimized amplification model.

**Supplementary Figure S2:**
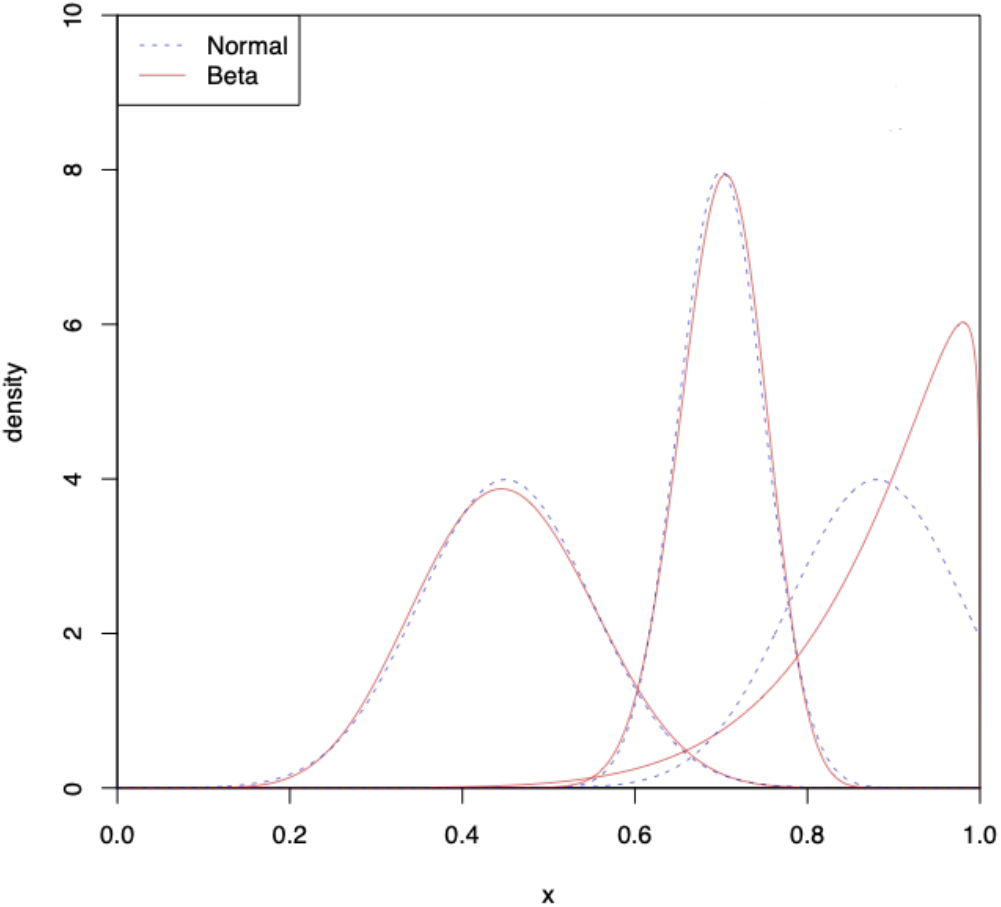
Comparison between the truncated normal and beta distributions for the inherited efficiency amplification model. The inherited efficiency model of Best et al. (2015) originally draws the replication efficiency for each clone (between 0 and 1) from a truncated normal distribution. This parametrization can be improved by drawing from a beta distribution instead. This makes very little difference within the range of intermediate values that Best *et al.* focus on (*e.g.*, means and standard deviations of 0.45±0.1 or 0.7±0.05, left and middle line pairs), but it simplifies and stabilizes the specification of the distribution when more extreme parameter values are used (*e.g.*, 0.9±0.1, right), because truncation becomes pronounced.

**Supplementary Figure S3:**
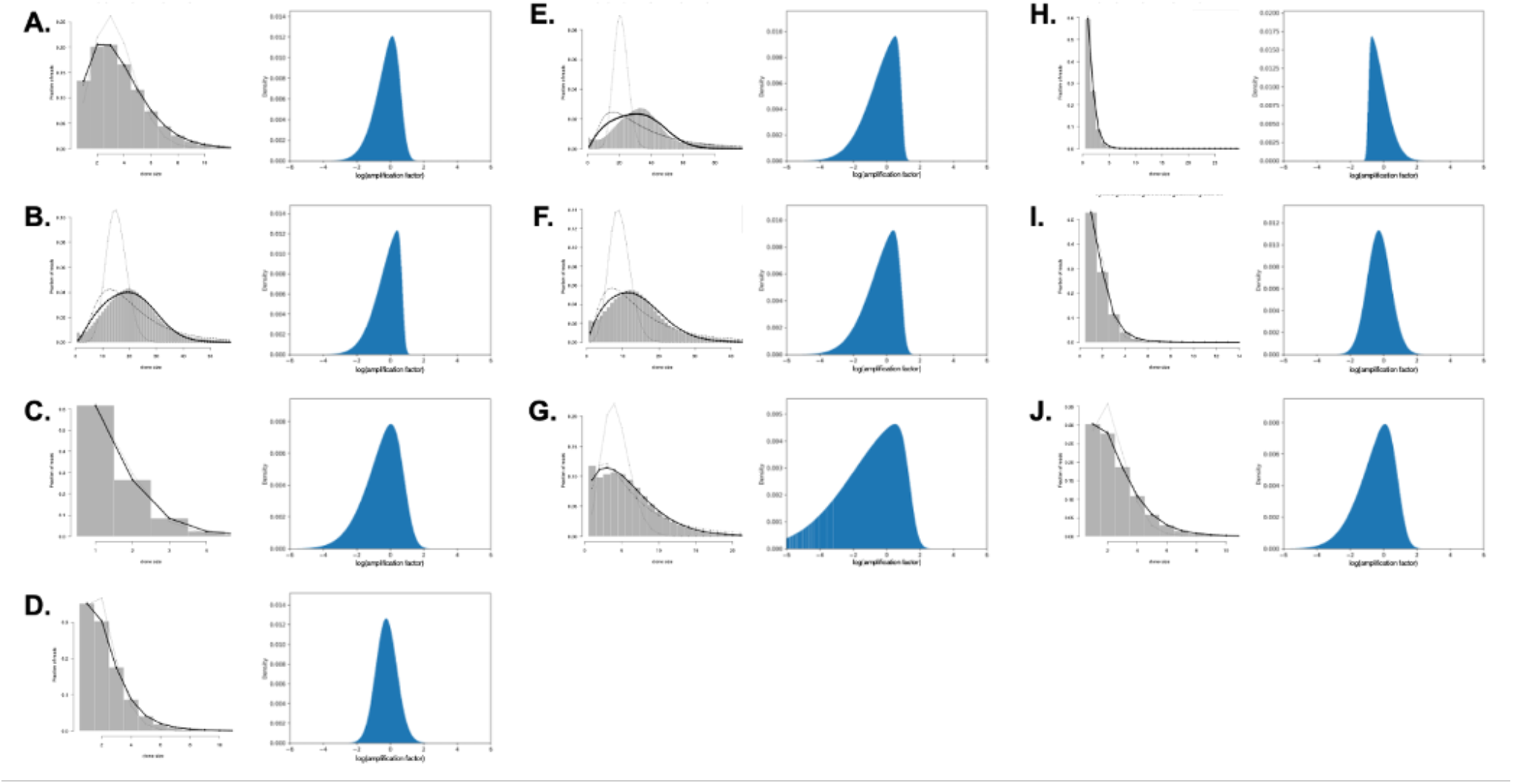
Fitted amplification models & depth-complexity ratios. Model fits to the read clone sizes histograms of all re-analyzed RAD libraries, namely (A) Anole-600ng, (B) Anole-30ng, (C) Robin, (D) Stickleback, (E) Penguin-3, (F) Penguin-4, (G) Penguin-81, (H) Warbler-1, (I) Warbler-2, and (J) Warbler-3. For each histogram, the solid line corresponds to a fitted depth-complexity ratio and log-skew-normal amplification noise model (itself shown on the right of the clone size histogram), the thin dashed line to a lognormal model, and the dotted gray line to a noiseless amplification model.it

**Supplementary Figure S4:**
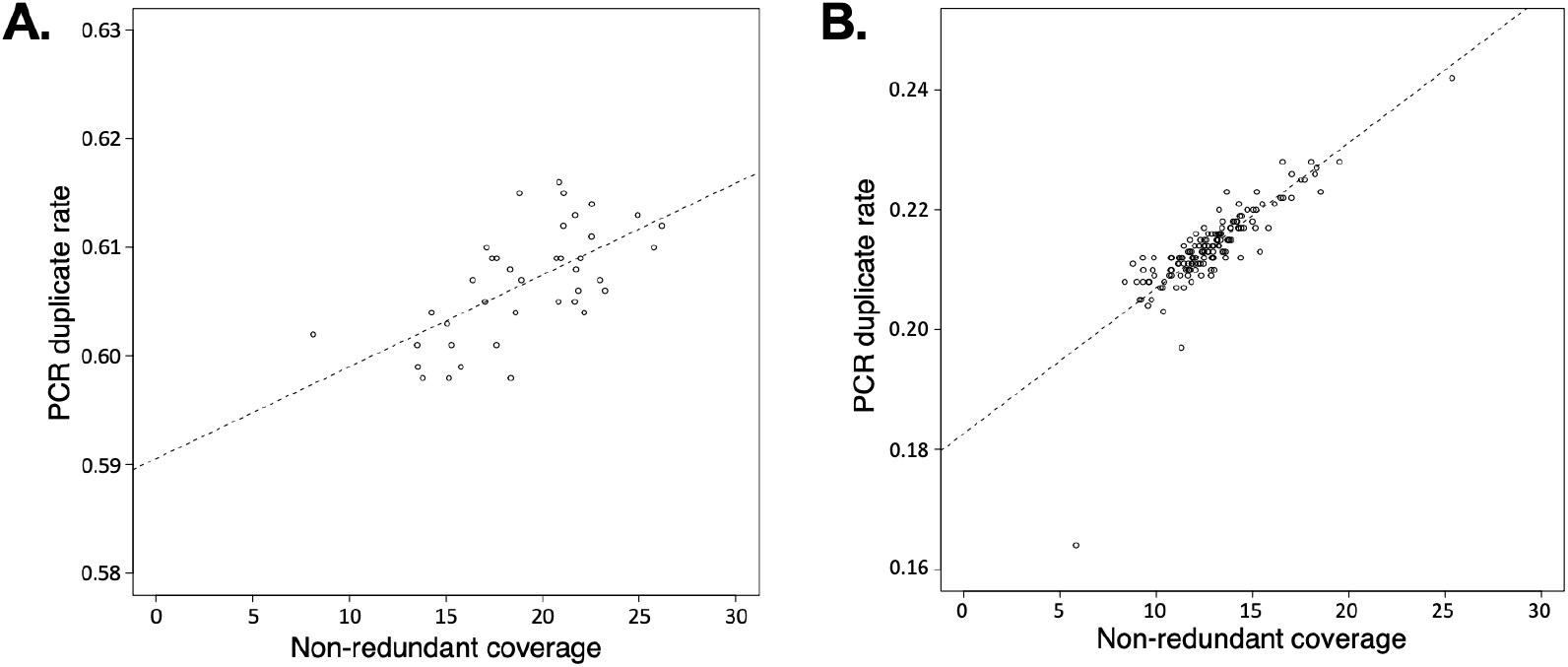
Within-library duplicate rate variation correlates with non-redundant coverage. Although PCR duplicate rates tend to be consistent among samples within a library, there is residual variation. Remarkably, this residual variation correlates with coverage for both the Anole-600ng (A; R^2^=0.41) and the Robin (B; R^2^=0.71).

**Supplementary Figure S5:**
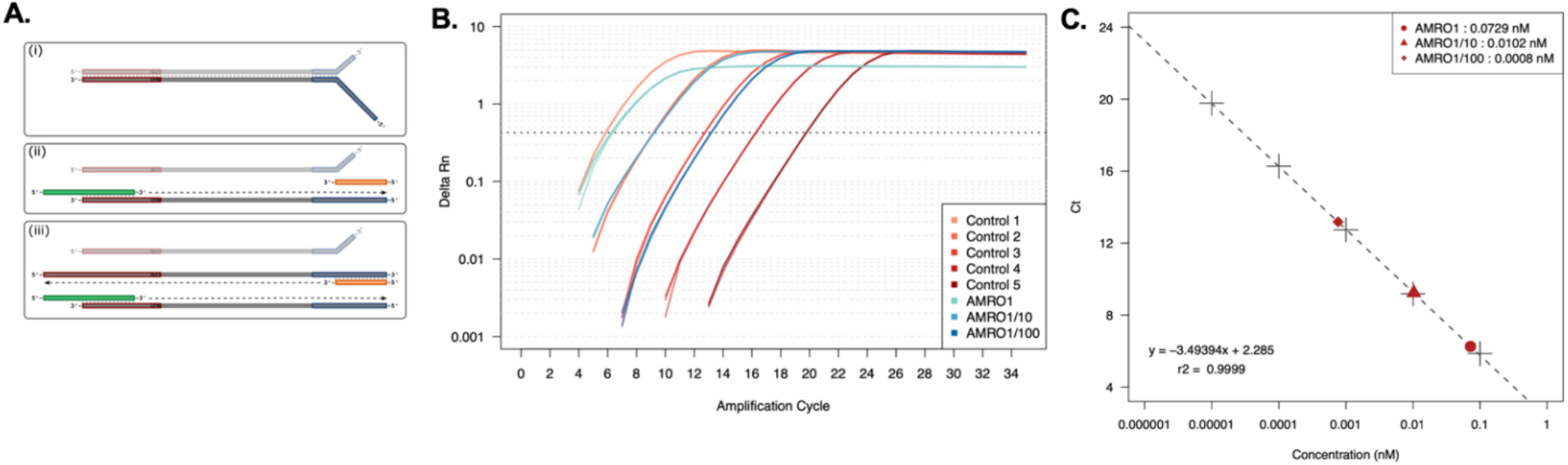
qPCR analyses. (A) Primer design and amplification mechanism (initial template structure (i); first cycle (ii); and second cycle (iii)) for RAD-seq libraries prepared following Baird et al. (2007), using the custom P1 adapters from (Hohenlohe et al., 2012). As such libraries rely on Y-adapters for selective amplification, for all intents and purposes they are effectively single-stranded. (B) Fluorescence curves for triplicated control and library dilution assays. (C) Measured concentrations for library dilutions in (B).

**Supplementary Figure S6:**
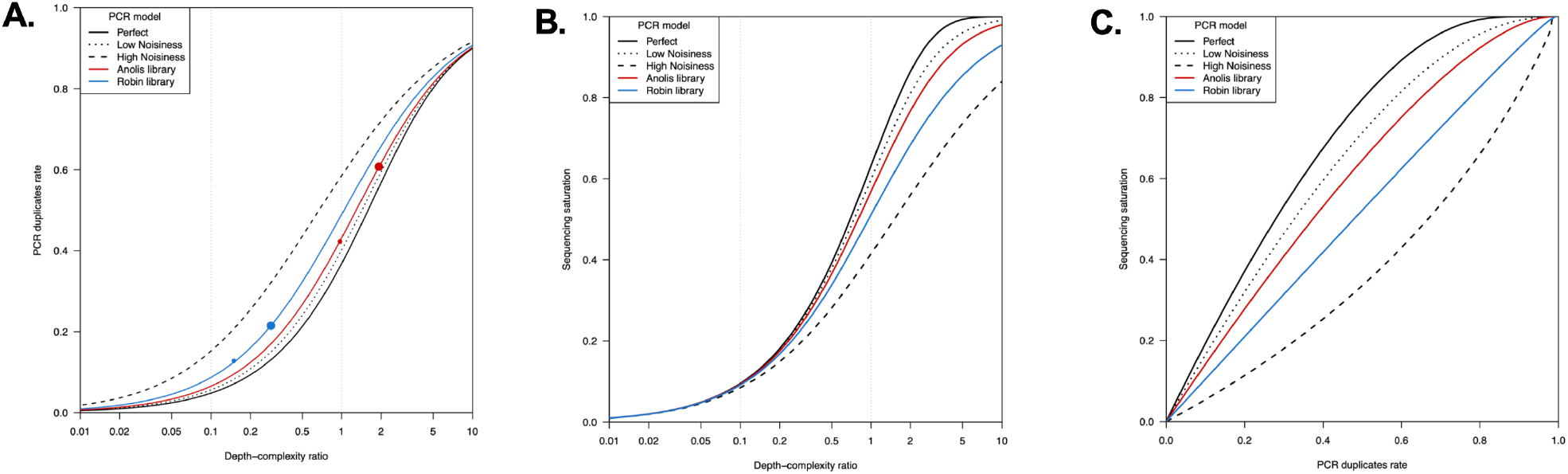
Depth-complexity ratio, PCR duplicate rate & sequencing saturation. The data shown in main text Figure 3C (reproduced here as panel A) can alternatively be used to illustrate how much sequencing saturation (*i.e.*, the fraction of clones sequenced at least once) is achieved at a particular depth-complexity ratio, for a range of amplification models (panel B), or to display the relationship between the PCR duplicate rate and the sequencing saturation (panel C).

